# Polymorphic KIR3DL3 expression modulates tissue-resident and innate-like T cells

**DOI:** 10.1101/2022.08.17.503789

**Authors:** William H. Palmer, Laura Ann Leaton, Ana Campos Codo, Patrick S. Hume, Bergren Crute, Matthew Stone, Adrie van Bokhoven, Richard P. Tobin, Martin D. McCarter, William J. Janssen, James Roest, Shiying Zhu, Jan Petersen, Julian P. Vivian, Jamie Rossjohn, John Trowsdale, Andrew Getahun, John Cambier, Liyen Loh, Paul J. Norman

## Abstract

Most human killer cell immunoglobulin-like receptors (KIR) are expressed by Natural Killer (NK) cells and recognize HLA class I molecules as ligands. Uniquely, KIR3DL3 is a conserved but polymorphic inhibitory KIR recognizing a B7 family ligand, HHLA2, and is implicated for immune checkpoint targeting. Because the expression profile and biological function of KIR3DL3 remained elusive, we searched extensively for KIR3DL3 transcripts, revealing expression is highly enriched in γδ and CD8+ T cells rather than NK cells. These KIR3DL3 expressing cells are rare in the blood and thymus, but more common in the lungs and digestive tract. High resolution flow cytometry and single cell transcriptomics showed that peripheral blood KIR3DL3+ T cells have an activated transitional memory phenotype and are hypofunctional. The TCR usage is biased towards genes from early rearranged TCR-α variable segments or Vδ1 chains. Whereas we detected no impact of KIR3DL3 polymorphism on ligand binding, variants in the proximal promoter and at residue 86 can reduce expression. Together, we demonstrate that KIR3DL3 is upregulated in response to unconventional T cell stimulation and that individuals may vary in their ability to express KIR3DL3. These results have implications for the personalized targeting of KIR3DL3/HHLA2 checkpoint inhibition.

## Introduction

Detection and elimination of infected or cancerous cells by cytotoxic lymphocytes is vital for survival ^1–3^. In this role, alongside those in placentation and tissue transplantation ^4–8^, killer cell immunoglobulin-like receptors (KIR) modulate the cytotoxic and inflammatory activity of human natural killer (NK) and T cells ^9,10^. Inhibitory KIR educate developing NK cells, enabling them to kill malignant or infected cells marked by altered cell surface ligand expression, namely HLA class I molecules ^9,10^. Expression of inhibitory KIR by T cells can also promote cell survival, dampen responses to T cell receptor stimuli, and suppress other auto-reactive T cells ^11–16^.

Among the 13 functionally distinct KIR molecules ^17–19^, KIR3DL3 is unusual. Whereas most *KIR* genes vary by presence and copy number ^17^, *KIR3DL3* is uniquely present in every individual across diverse populations ^20^ and conserved among humans and catarrhine primates ^21^. *KIR3DL3* is also the most heterozygous *KIR* ^22^, with a subset of polymorphic residues having genetic patterns consistent with the action of diversifying natural selection ^21^. Recently, from high-throughput *in vitro* fusion protein microarray screening, KIR3DL3 was found to recognize the B7 family protein HHLA2 as a ligand ^23–26^, in contrast with other KIR that recognize HLA class I as ligands. HHLA2 expression by target cells coordinates inhibition of cytotoxic responses by KIR3DL3-transduced NK cell lines ^23,24^, consistent with the presence of an immunoreceptor tyrosine-based inhibitory motif (ITIM) in the KIR3DL3 cytoplasmic tail ^21^. These evolutionary patterns and ligand binding restriction suggest an essential immunoregulatory function of KIR3DL3. However, the immunological role of KIR3DL3 remains unclear.

The lack of a described expression pattern for KIR3DL3 has been a considerable obstacle to understanding the biological importance of this inhibitory receptor. Most KIR are expressed stochastically and frequently by peripheral NK cells ^27–29^, but efforts to localize KIR3DL3 on peripheral NK cells have produced contradictory results. KIR3DL3 expression in peripheral blood, as assayed by PCR, was found to be undetectable in 80% of donors ^30^, and perhaps enriched in CD56^bright^ peripheral blood and decidual NK cells ^31^. However, more recently, widespread KIR3DL3 expression was described in peripheral CD56^dim^ NK cells ^24^, while another report showed induced cell surface expression in peripheral blood CD4+ and CD8+ T cells following *in vitro* TCR-mediated activation and long-term cell culture ^23^. Distinct transcription factor binding sites in the KIR3DL3 promoter support a unique expression profile as compared to other KIR ^32,33^. Altogether, the presently available data describing KIR3DL3 expression are incongruent, fragmented, and warrant a wide-ranging search for KIR3DL3 across tissues and cell types.

KIR3DL3 likely represents an immune checkpoint, with the HHLA2-KIR3DL3 interaction suggested as an immunotherapeutic target. Other B7 family ligands and their receptors, such as PD-L1 and CTLA-4, have been the targets for successful immunotherapies ^34,35^. Indeed, HHLA2 is upregulated in tumors of the lung, gastrointestinal tract, kidney, and liver, and in some cases, expression has been associated with poor prognosis ^36–39^. Given KIR3DL3 is an inhibitory receptor, expression on tumor infiltrating lymphocytes could lead to immune evasion by HHLA2+ tumors. Prerequisite to harnessing any medicinal utility is a basic characterization of KIR3DL3 expression and the function of its polymorphism. Given the high prevalence of functional polymorphism in other KIR proteins ^40–48^ and promoters ^49,50^, a consideration of KIR3DL3 diversity is paramount to understanding the role of KIR3DL3 across human populations. In this study, using a highly specific monoclonal antibody for KIR3DL3 in combination with multi-dimensional spectral flow cytometry and single cell transcriptomics, we decipher the tissue distribution of KIR3DL3, and its immunoregulatory function. Further, we determine functional consequences of selected promoter and amino acid polymorphism. These studies shed fundamental biological insight into this enigmatic KIR family member.

## Results

### KIR3DL3 RNA expression is found in cytotoxic lymphocytes from diverse tissues

Although *KIR3DL3* expression is most often silenced through promoter methylation, *KIR3DL3* mRNA has been identified previously in CD56^bright^ peripheral blood and decidual NK cells ^30,31^. To better determine the extent of *KIR3DL3* transcription, we examined RNA sequencing experiments publicly available from the NCBI short read archive (SRA). By searching for a unique *KIR3DL3*-characterizing sequence (see Methods), we identified KIR3DL3 mRNA in 196 of the 360,787 available data sets. After excluding cell lines, *ex vivo* experiments and replicated individuals, we ultimately identified 84 individuals across 48 studies showing evidence for KIR3DL3 expression (Table S1). Of the 84 individuals, we did not observe any evidence for differences in expression levels (as determined by normalized read counts) of KIR3DL3 across ages, between males and females, between donors who were healthy or with disease, nor any interactions between age, sex, and health status (Table S1). However, 7 of 9 KIR3DL3+ samples from the lung, and an additional lung-draining lymph node sample (Table S1), were from individuals with non-small cell lung cancer, which is often marked by high HHLA2 expression ^37^. Thus, while we did not observe a general effect of health status on *KIR3DL3* mRNA expression, expression may still be enriched in specific malignant or immunopathological diseases.

Cell sorted cytotoxic lymphocytes derived from diverse tissues constituted the majority of samples positive for *KIR3DL3* mRNA (Fig. 1A; Table S1). Among these tissues, *KIR3DL3* transcripts were found most often in the digestive tract, lung, and lymphoid organs, with each of these systems being represented by multiple independent studies (digestive: 5; lung: 6; lymphoid: 4). We also identified KIR3DL3 transcripts from decidual T cells, consistent with previous reports of KIR3DL3 expression by CD56^bright^ decidual NK cells ^31^. We observed sequence read coverage across all KIR3DL3 exons (Fig. 1B). Remarkably, the tissues identified as most likely to express KIR3DL3 (Fig. 1A,C) overlap considerably with HHLA2 protein expression patterns ^51^ (Fig. 1D), suggesting either KIR3DL3+ cells are recruited to these tissues or KIR3DL3 is induced in ligand-rich microenvironments. In summary, we detected KIR3DL3 mRNA in peripheral blood mononuclear cells (PBMCs) and tissues from the respiratory, digestive, and lymphoid systems. We next sought to confirm that presence of full KIR3DL3 transcripts correlates with cell surface protein expression.

**Figure 1:**
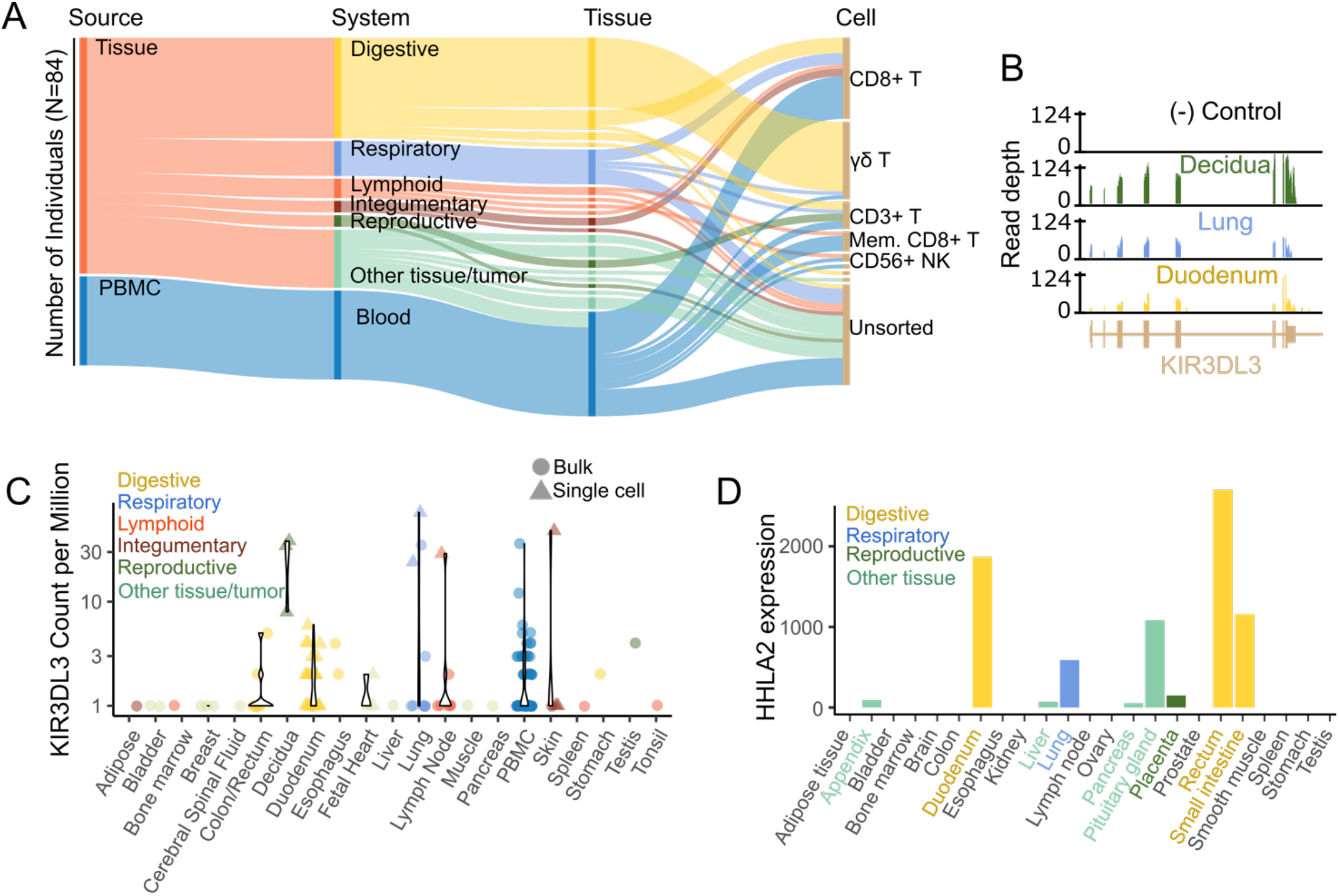
KIR3DL3 mRNA identified in PBMCs and tissues that express HHLA2. (A) The first million reads from each of 360,787 short read archive (SRA) records were searched for a unique and specific KIR3DL3 subsequence. Shown are the frequency of tissue source, systems, tissues, and sorted cell types of the 84 individuals so identified as having KIR3DL3 transcripts. (B) All sequence reads from three KIR3DL3+ samples were mapped to the hg38 human genome reference. Each sample showed high depth of coverage across KIR3DL3 exons. (C) Plotted is the frequency of the KIR3DL3 subsequence per million reads. Colors indicate tissue system, symbols indicate bulk (circles) or single-cell (triangles) sequencing experiments. (D) HHLA2 protein expression as described by *Wang et al* ^51^.

### Cell surface KIR3DL3 is more frequently found in lung and the digestive tract tissues

Guided by the SRA data, we used multi-parameter spectral flow cytometry to delineate the specific lymphocyte subsets that express cell surface KIR3DL3 protein. Thus, we used single cell suspensions of PBMCs, lung, colon, and thymus tissues. The CH21 antibody specifically stains KIR3DL3 ^31^ (Fig. S1 for antibody specificity) (Fig. S2 for gating strategy), and consistently identified a small yet distinct population of non-B cell lymphocytes among PBMCs (Median= 0.007%, n=13, p < 0.001, Fig. 2A-B, Fig. S3). In thymic tissues, KIR3DL3 expressing cells were found at a similar frequency as observed in peripheral blood (Median=0.006%, n=3, Fig. 2A-B). Consistent with SRA data (Fig. 1A-B), we found that the proportion of KIR3DL3+ cells was significantly higher in lung lymphocytes (0.04% of non-B lymphocytes; n=5, p = 0.03), colon intra-epithelial lymphocytes (IELs) (0.06%; n=3, p = 0.04), and small intestine IELs (0.45%; n=3 p < 0.001) as compared to peripheral blood (Fig. 2B). We did not observe any correlation between the frequency of KIR3DL3+ cells with either sex (p = 0.49) or age (p = 0.16). These results confirm that SRA data reflects the protein expression of KIR3DL3, which occurs more frequently in lymphocytes of mucosal tissues.

**Figure 2:**
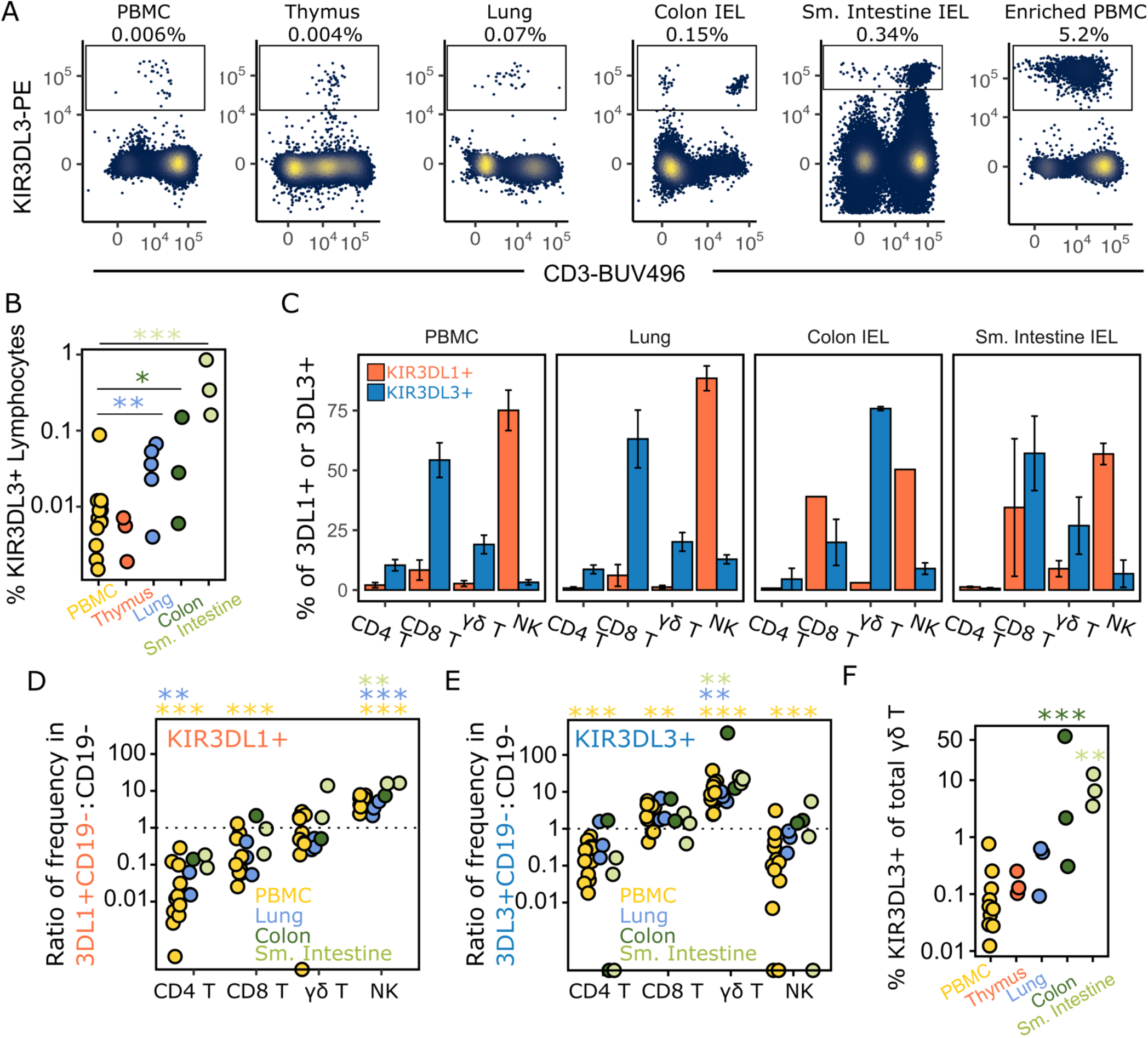
KIR3DL3 is enriched in CD8+ and γδ T cells of PBMC, lung, and digestive tract IELs. Human PBMCs, thymocytes, lung lymphocytes, and digestive tract intra-epithelial lymphocytes (IELs) were analyzed using 17-color spectral flow cytometry. (A) Shown is the percentage of cells expressing KIR3DL3 in the viable CD14-CD19-population, isolated from the tissue indicated at the top each panel. (B) Summary of (A) across ten PBMC, three thymus, five lung, three colon, and three small intestine donors. (C-E) Magnetic bead enrichment (e.g. Enriched PBMC) was used to increase the number of KIR3DL3+ cells prior to flow cytometry. (C) The percentage of CD4+ T, CD8+ T, γδ T, and NK cells was calculated for KIR3DL1+CD19- and KIR3DL3+CD19-lymphocytes. These frequencies (C) were compared to the background (CD19-) to determine the enrichment of KIR3DL1 and KIR3DL3 expression in CD4+ T, CD8+ T, γδ T, and NK cells (D-E). Significant differences in the frequency of lymphocyte subsets within CD19- and KIR+CD19-parent populations are indicated, with asterisks colored by tissue. (F) Shows the proportion of γδ T cells that express KIR3DL3 according to the tissue of origin.

### KIR3DL3 is expressed in CD8+ and γδ T cells of peripheral blood and tissues

To more specifically characterize the lymphocyte subsets that express KIR3DL3, we enriched for KIR3DL3+ cells with a magnetic bead-based enrichment using anti-PE microbeads bound to CH21 and secondary anti-Ms-PE (Fig. 2A). We then calculated NK and T cell subset frequencies in all CD19- (i.e. background), CD19-KIR3DL3+, or CD19-KIR3DL1 + lymphocytes (Fig. 2C) and determined enrichment of these T/NK cell subsets in either KIR3DL1+ (Fig. 2D) or KIR3DL3+ (Fig. 2E) cells, as compared to the background frequency. Both KIR3DL1 and KIR3DL3 were significantly less likely to be expressed by CD4+ T cells (p<0.001) and KIR3DL1 expression was highly enriched on NK cells (p < 0.001), as expected (Fig 2D-E). Strikingly, across PBMC and tissues, KIR3DL3 was most enriched on γδ T cells (p <0.001), followed by CD8+ T cells (p = 0.002) (Fig. 2E). In PBMCs, while the median proportion of γδ T cells of CD19-peripheral lymphocytes was 2.1%, 16.3% of KIR3DL3+CD19-cell lymphocytes were γδ T cells. KIR3DL3+ lymphocytes of the lung, small intestine, and colon mirrored these patterns, with KIR3DL3 expression enriched in lung CD8+ T cells (lung: p = 0.06) and in lung, colon, and small intestine γδ T cells (lung: p = 0.002; colon: p = 0.08; sm. Intestine: p = 0.006). The absolute frequency of KIR3DL3 expression was most striking in digestive tract IELs, where an average of 14% (range: 0.3%-58%) of γδ T cells expressed KIR3DL3 (Fig. 2F).

Previous studies have shown that KIR can be expressed on the cell surface of terminally differentiated CD8+ T cells ^11^. To determine the differentiation and memory phenotypes of KIR3DL3+ T cells across the tissues, we analyzed the cell surface markers CD27 and CD45RA in combination to derive subsets of naïve-like (CD27+CD45RA+), central memory-like (CM, CD27+CD45RA-), effector memory-like (EM, CD27-CD45RA-), and EM re-expressing CD45RA (EMRA, CD27-CD45RA+)-like T cells. As above, we compared background CD8+ or γδ T cells to KIR3DL1+ or KIR3DL3+ T cells to determine if these KIR are preferentially expressed in certain T cell subsets. Most notably, we observed preferential KIR3DL3 expression on CD27^hi^ cells, with greater CD27 mean fluorescent intensity (MFI) as compared to background naïve or CM populations (Fig. 3A). Interestingly, KIR3DL3+ γδ T cells of peripheral blood and lung were significantly more likely to co-express CD27 and CD45RA (PBMC, p = 0.014; lung, p = 0.046), suggestive of a naïve-like phenotype (Fig. 3A-B; Fig. S4). KIR3DL3+ CD8+ T cells were distributed across all four differentiation states, with significantly less EM-like subsets in PBMCs (Fig. 3B, p=0.042). Conversely, KIR3DL1 expression on PBMCs was preferentially enriched in CD8+ EMRA T cells (Fig. 3B, p = 0.014), although this enrichment did not exist in γδ subsets (Fig. 3B, p = 0.52). Consistent with the mature nature of IELs, we observed that almost all colon and small intestine IELs were EM or EMRA-like, including KIR3DL3+ cells (Fig. 3B).

**Figure 3:**
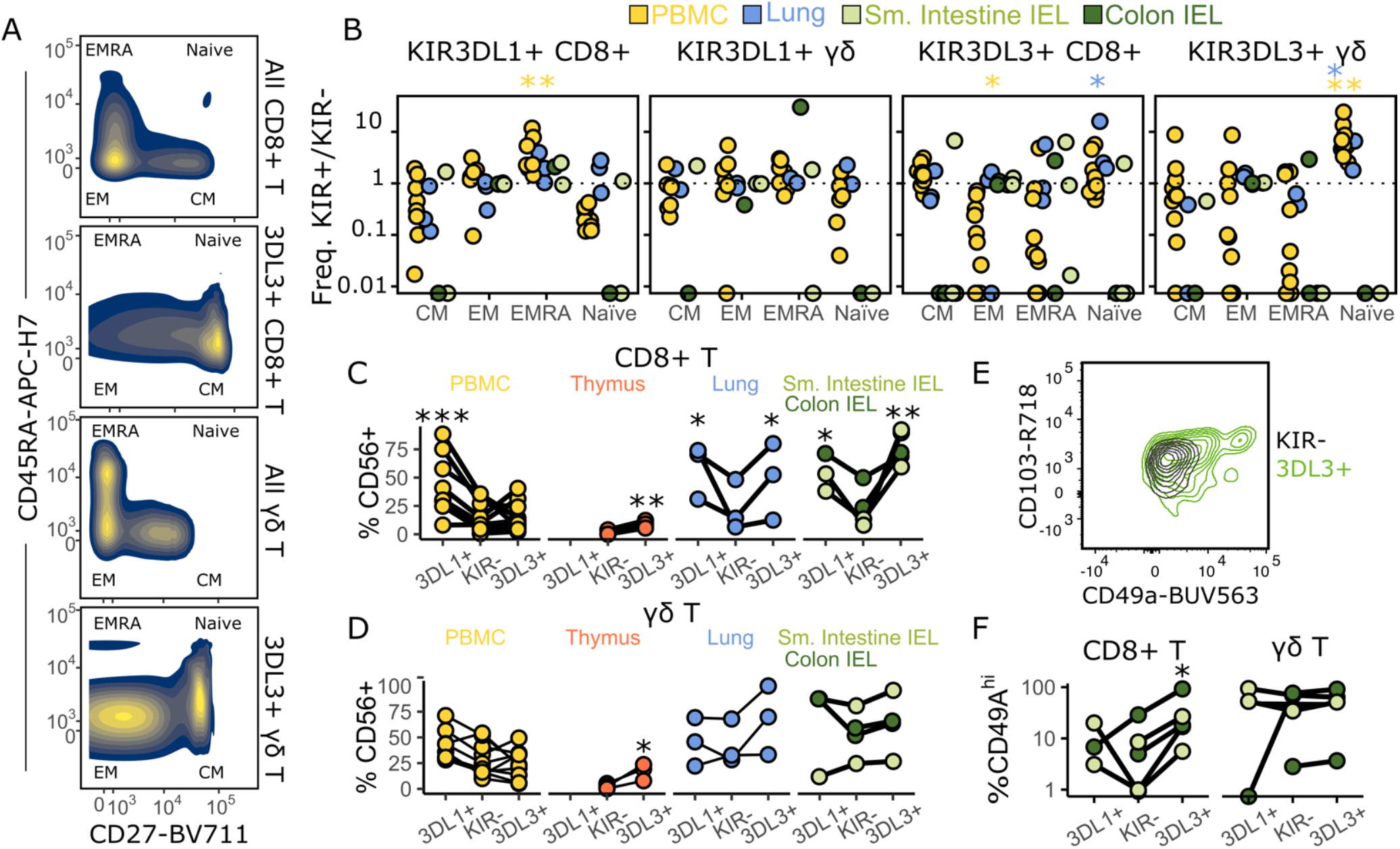
KIR3DL3+ T cells can co-express naïve, innate, and tissue-resident T cell markers. (A) Distribution of CD27 and CD45RA on CD8+ T or γδ T cells unselected (upper) or selected (lower) for KIR3DL3 expression. Average kernel density across five PBMC donors is shown. EM=effector memory, EMRA=EM re-expressing CD45RA, CM=central memory. (B) Frequency of CM-, EM-, EMRA-, and naïve-like CD8+ or γδ T cell populations in KIR3DL1 + or KIR3DL3+ cells were divided by the frequency of these populations in KIR3DL1- or KIR3DL3-cells. (C) The proportion of cells expressing CD56 on KIR-, KIR3DL1+, or KIR3DL3+ CD8+ T or (D) γδ T cells is plotted. Each line represents one donor. (E) Representative contour plot of CD49a^hi^ population on KIR3DL3+ CD8+ T cells from the intestine. (F) The summarized proportion of CD49hi cells of 3DL1+, KIR-, or 3DL3+ CD8+ and γδ T cells. Significance assessed by linear mixed models. *p<0.05, **p<0.01, ***p<0.001

Analyses of tissue residency markers and CD56, constitutively expressed on NK cells and populations of cytotoxic and innate-like T cells ^52^, revealed an altered expression of CD56 and CD49a on KIR3DL3+ T cells. KIR3DL3+ CD8+ T cells had greater expression of CD56 in the thymus (p=0.003), lungs (p=0.031), and digestive tract (p=0.001), compared to KIR-cells (Fig. 3C). Although KIR3DL3+ γδ T cells had similar trends in CD56 expression to KIR3DL3+ CD8+ T cells, significant upregulation was only observed in the thymus (Fig. 3D, p=0.03). In comparison, KIR3DL1+ CD8+ T cells, had higher CD56+ than KIR-cells in PBMCs (p<0.001), in addition to all other tissues (lung, p = 0.01; IEL, p=0.05), with the exception of thymus, where there was no detectable expression of KIR3DL1. In most tissues, we did not observe differential expression of tissue residency markers CD103 and CD49a in KIR3DL3+ or KIR3DL1+ cells. The exception was in IELs, where a subset of KIR3DL3+ CD8+ T cells had a CD49a^hi^ phenotype (Fig. E-F, p=0.03). In summary, KIR3DL3 is enriched in CD8+ T and γδ T cells across tissues, is co-expressed with CD27 in PBMC and lungs, and in more mature (i.e. CD27-), tissueresident T cell subsets in digestive tract IELs.

### KIR3DL3+ T cells have an altered effector cell transcriptional profile

To characterize the determinants of KIR3DL3 expression to higher resolution, we performed scRNA sequencing on sorted populations of KIR3DL3+ PBMCs, alongside NK, CD8+ T, and γδ T cells, from five donors. We obtained transcriptomic data for 9,188 NK, 2,750 CD8+ T, 3,490 γδ T, and 1,050 KIR3DL3+ cells. Using Seurat, ^53^ we identified 9 distinct NK and T cell clusters (Fig. 4A), with KIR3DL3+ cells corresponding to a unique subpopulation of cells (Fig. 4B). Multimodal reference mapping of gene expression profiles to a CITE-seq dataset of immune cells from peripheral blood ^53^ suggested KIR3DL3+ cells have a transcriptional profile intermediate between canonical naïve and memory cell populations (Fig. 4C). KIR3DL3+ cells were nominally defined as EM CD8+ T cells, converse to their designation as naïve-like or CM-like by cell surface staining for CD27 and CD45RA (Fig. 3B). To quantify the naivety of KIR3DL3+ cells, we used VISION ^54^ to calculate a Naïve T cell Score, using markers that defined naïve T cells in our data. Indeed, we observed that KIR3DL3+ T cells were more transcriptionally naïve than background EM or γδ T cell subsets (p<0.001; Fig. 4D). Differential gene expression analysis (Table S2) revealed that this intermediate phenotype was due to the simultaneously low expression of genes encoding cytotoxic granule proteins (*PRF1, NKG7, GZMA, GZMB, GZMH, FGFBP2*) and those involved in protein translation (*EEF1A1*) and ribosomal biogenesis (*RPS8, RPS6, RPL34, RPL39, RPL31, RPS4X, RPS13, RPL3*) (Fig. 4E). Intracellular staining of PBMCs and analysis by flow cytometry agreed with the transcriptional profile, where reduced GzmB, GzmK, and Perforin protein expression was found in KIR3DL3+ CD8+ T and γδ T cells compared to total EM, CM or γδ T cells (Fig. 4F). Low expression of ribosome biogenesis genes is correlated with poor T cell proliferative capacity ^55^. To confirm KIR3DL3+ T cells have an altered proliferative capacity, we stimulated T cells in PBMC (n=4 donors) using magnetic beads covalently labeled with anti-CD3 and anti-CD28 to engage the TCR complex. Following 7 days, the absolute number of stimulated T cells expanded by 3.7-fold as compared to non-stimulated controls (Fig. 4G). Conversely, the number of KIR3DL3+ T cells remained unchanged (p < 0.001) (Fig. 4G), suggesting that these cells did not proliferate. As such, KIR3DL3+ T cells are an anomaly along the ‘innateness’ gradient ^55^, which proposes the gradual gain of effector function concomitant with the loss of proliferative capacity through reduced ribosome biogenesis. Instead, KIR3DL3 expression may mark a hyporesponsive T cell population with limited naïve-like effector function, but an effector-like proliferative capacity.

**Figure 4:**
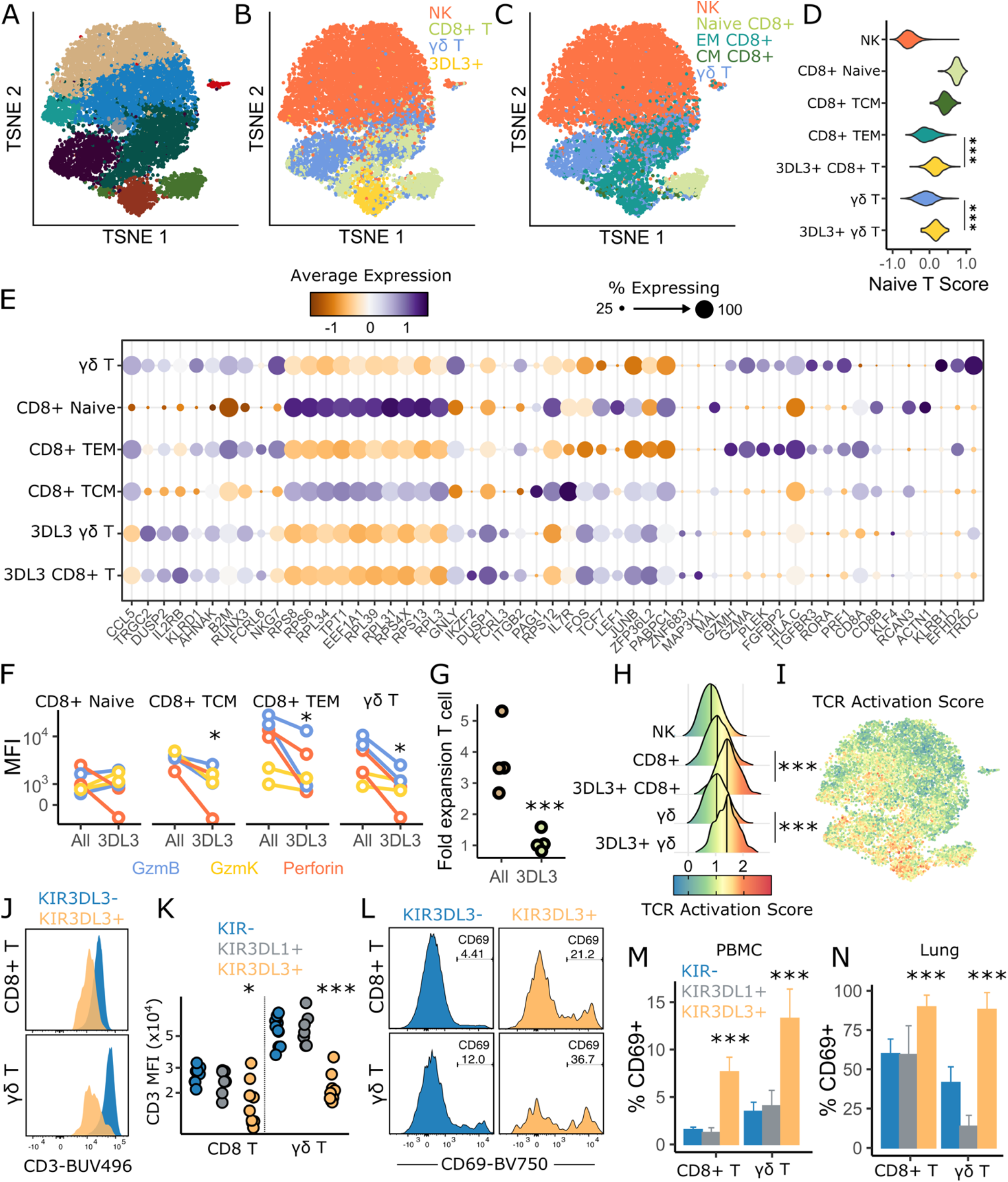
KIR3DL3+ cells are hypofunctional with a signature of recent TCR activation. (A-C) Single cell RNA sequencing of sorted NK, CD8+ T, γδ T, and KIR3DL3+ cells from PBMC of 5 donors; visualized using TSNE, colored according to Seurat cluster (A), sorted population (B), or by T cell subpopulation as assigned by multimodal reference mapping (C). (D) VISION was used to calculate a Naïve T Score, using genes that defined CD8+ naïve T cells in our data. (E) KIR3DL3+ cells in each T cell subpopulation (from panel (C)) were compared to respective KIR3DL3-populations for differential gene expression testing in Seurat. Dot plots show the percentage of cells expressing each gene named underneath and the average scaled expression within each subpopulation. (F) PBMC from 2 donors were stained for GzmB, GzmK, and Perforin and mean fluorescence intensity (MFI) determined by flow cytometry. (G) PBMC from 4 donors were stimulated with T Cell-Activator dynabeads for a week and fold expansion of either all T cells or KIR3DL3+ T cells was calculated as relative to unstimulated T cells. (H-I) VISION was used to calculate a T Cell Activation score (Fig. S5 for gene list), plotted as the distribution of the score of each cell across cell subsets (H) or by color on the TSNE dimension reduction (I). (J) Representative histograms of CD3 expression on KIR3DL3- (blue) or KIR3DL3+ (orange) CD8+ T cells (upper panel) and γδ T cells (lower panel) from PBMCs. (K) Mean fluorescence intensity of CD3 expression on KIR- (blue), KIR3DL1+ (grey) and KIR3DL3+ (orange) CD8+ T cells or γδ T cells, n= 10 PBMC donors. (L) Representative histograms showing the proportion of CD69+ KIR3DL3- (blue) or KIR3DL3+ (orange) on CD8+ T cells (upper panel) and γδ T cells (lower panel) from PBMCs and summarized in PBMC (M, n=10) and lungs (N, n=3). Significance assessed by linear mixed models. *p<0.05, **p<0.01, ***p<0.001

### TCR-responsive genes define KIR3DL3+ T cells

The most significantly upregulated gene among KIR3DL3+ CD8+ and γδ T cells was the TCR-inducible *FOS,* as compared to KIR3DL3-CD8+ (p = 2.5×10^-75^) and γδ T (p = 4.7xl0^-14^) cells, respectively (Fig. 4E, Table S2). Indeed, the most highly induced genes included many known to be upregulated following TCR stimulation (e.g. JUN, FOSB, IL2RB, DUSP2, EGR1) ^56,57^. To test the degree with which KIR3DL3+ cells were enriched for upregulation of TCR-responsive genes, we again used VISION to compute an integrated TCR Activation Score, using an unbiased set of established TCR-responsive genes ^56,57^ (Fig. S5A). We found that KIR3DL3+ CD8 and γδ T cells had a significantly greater TCR activation score as compared to their KIR3DL3-counterparts (Fig. 4H-I). Consistent with recent TCR stimulation, we also observed reduced CD3 MFI on KIR3DL3+ T cells (Fig. 4J-K). Analyses of the recent activation and tissue resident memory marker, CD69, in PBMC and lung lymphocytes revealed that both KIR3DL3+ CD8+ (PBMC, p<0.001; lung, p<0.001) and γδ (PBMC, p<0.001 ; lung, p<0.001) T cells expressed higher proportions of CD69+ compared to their KIR3DL3-counterparts (Fig. 4L-N). Differences in either CD3 MFI or frequency of CD69+ cells were not observed in KIR3DL1+ T cells (Fig. 4K,M-N). The majority of digestive tract IELs were CD69+, with no differences across KIR- or KIR+ T cells, consistent with a tissue resident profile. KIR3DL3+ thymocytes also were more likely to express CD69, CD27, and be CD4+CD8+, markers consistent with recent positive thymic selection (Fig. S5B-C).

Given the hyporesponsive transcriptional profile of KIR3DL3+ T cells, their lack of proliferation in response to TCR-mediated stimulation and reduced cytolytic molecules, we hypothesized KIR3DL3 is expressed as an immune checkpoint in auto-reactive T cells. Such cells may experience agonistic selection or chronic T cell stimulation. Gene set enrichment analyses ^58^ identified NF-κB targets as the most significantly upregulated gene set in KIR3DL3+ cells (CD8 p^adj^ = 0.002; γδ p^adj^ = 0.02), alongside downregulated metabolism genes such as those involved in oxidative phosphorylation (CD8 p^adj^ = 0.006; γδ p^adj^ = 0.17) or mTOR targets (CD8 p^adj^ = 0.08; γδ p^adj^ = 0.03). Both NF-κB and mTOR are required for sustained T cell activation ^59^ and oxidative phosphorylation can be reduced in T cells exhausted by chronic antigen stimulation ^60^. In summary, the transcriptional profile of KIR3DL3+ T cells is consistent with recent TCR stimulation, but without progression into an effector program.

### KIR3DL3 is enriched on Vδ1+ γδ T cells and CD8+ T cells with early-rearranging TCR-α chains

KIR3DL3 is expressed by both γδ and CD8+ T cells (Fig. 2), although transcriptionally, they are distinct from conventional naïve and effector/memory T cell subsets (Fig. 4B). To understand whether the TCR repertoire of KIR3DL3+ T cells is clonally restricted or diverse, we performed single cell paired VDJ sequencing of the α and β or γ and δ TCR chains. We obtained paired TCR chain sequences from a total of 1,695 CD8+ αβ T cells, 1,252 γδ T cells, 399 KIR3DL3+ CD8+ αβ T cells, and 66 KIR3DL3+ γδ T cells. Diverse αβ V gene pairing was observed in KIR3DL3+ CD8+ T cells, with CDR3 diversity similar between KIR3DL3+ and KIR3DL3-αβ T cells (α-chain, p=0.7; β-chain, p=0.8). Strikingly, we observed that KIR3DL3+ γδ repertoire was dramatically altered as compared to the total γδ TCR repertoire (Fig. 5B-C). Vγ9Vδ2 are the most common subset of γδ T cells in peripheral blood, generated during gestation ^61^. KIR3DL3 expression was identified only rarely on these cells, instead being preferentially expressed in Vδ1 and to a lesser extent, Vδ3 γδ T cells (Fig. 5C). To confirm Vδ usage across tissues, we stained KIR3DL3+ T cells from peripheral blood, lung, thymus, and intestinal IELs with Vδ1 and Vδ2 specific antibodies (Fig. S2). Indeed, for KIR3DL3+ γδ T cells, the proportion of Vδ1+ is enriched over Vδ2+ cells in peripheral blood, and lungs (Fig. 5D-F). KIR3DL3+ γδ T cells were almost exclusively Vδ2-, in contrast to KIR3DL1+ γδ T cells which expressed either Vδ1 or Vδ2 (Fig. 5E-F). Likewise, in the thymus KIR3DL3+ γδ T cells were also Vδ1+ with negligible Vδ2 expression (Fig. 5G).

**Figure 5:**
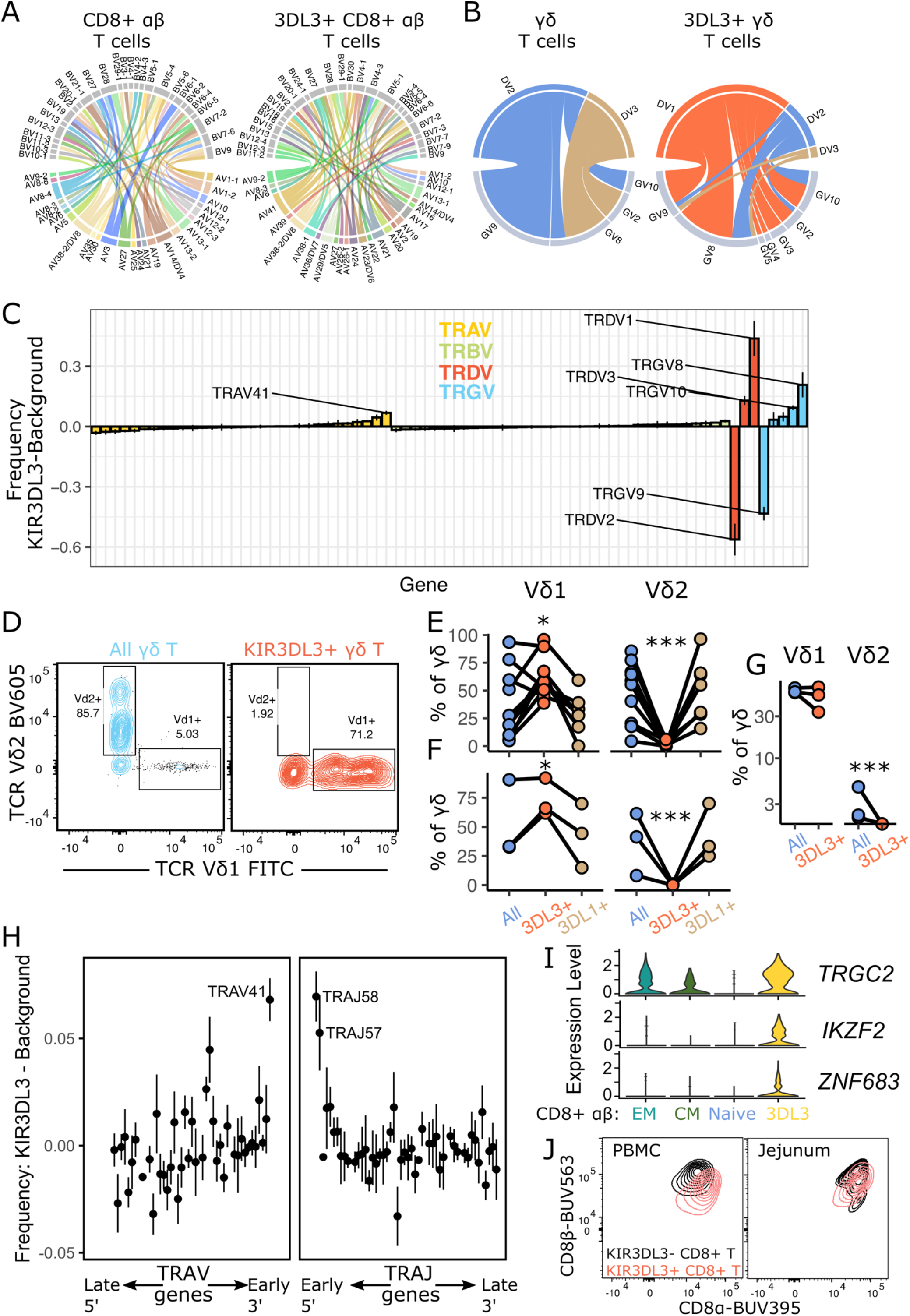
KIR3DL3 is preferentially expressed on T cells utilizing Vδ1 or early rearranged α chain TCRs. Single cell TCR sequencing was performed on PBMCs from 5 individuals by cell sorting KIR3DL3+, total γδ, or CD8+ αβ T cells. (A-B) Representative Circos plots of paired TCR variable (V) α (TRAV) and β (TRBV) chain usage, (A) or variable γ chain (TRGV) and variable δ chain (TRDV) pairings (B) from a single donor. Each segment represents paired TCRαβ or TCRγδ usage, and the thickness of each segment reflects the frequency of the paired usage; segments are colored according to the corresponding TRAV or TRDV usage. (C) V gene usage is summarized across 5 donors by subtracting the frequency of each V gene in KIR3DL3+ cells by that in KIR3DL3-cells. Variable genes significantly associated with KIR3DL3 expression are labelled. (D) Representative contour plots of PBMCs stained with fluorescently-conjugated Vδ1 and Vδ2 antibodies and analyzed by flow cytometry, the proportion of Vδ1+ and Vδ2+ cells are shown of total γδ T cells or KIR3DL3+ γδ T cells. Plotted are the proportions of Vδ1 and Vδ2 cells of total γδ, KIR3DL1+ γδ, and KIR3DL3+ γδ T cells across 9 PBMC (E), 3 lung (F), and 3 thymus donors (G). (H) Enrichment of TRAV and TRAJ usage in KIR3DL3+ T cells was plotted according to the propensity for each gene to undergo early TCR-α chain rearrangement (TRAV ranked 3’→ 5’; TRAJ 5’ → 3’). (I) Violin plots showing gene expression levels for: TCR gamma chain (*TRGC2*), Helios (*IKZF2*), and Hobit (*ZNF683*) CD8+ αβTCR effector memory (EM), central memory (CM), naive and 3DL3 clusters. (J) Peripheral blood and jejunum intra-epithelial CD8+ T cells were stained with fluorescent CD8α and CD8β antibodies and expression detected by flow cytometry. Significance assessed by linear mixed models. *p<0.05, **p<0.01, ***p<0.001

In the CD8+ KIR3DL3+ αβ T cells, when considering each TRAV or TRBV gene individually, TRAV41 was the only gene significantly enriched (Fig. 5C). Interestingly, TRAV41 gene usage defines an innate T cell population in some mammals ^62^ and in humans is located in the most proximal 3’ gene position of the TRAV region. The latter is noteworthy because innate-like CD8αα+ T cells enriched in intestinal tissues likely develop by thymic agonistic selection marked by early TCR-α gene re-arrangement ^63–66^. Early TCR-α rearrangement preferentially utilizes the TRAV and TRAJ genes that are closest in proximity, with TRAV41 in closest proximity to the TRAJ cluster. We therefore tested whether TRAV and TRAJ gene usage in KIR3DL3+ CD8+ T cells were biased towards early rearranging TCR-α chains. Indeed, we observed preferential usage for 3’ TRAV (p <0.001) and 5’ TRAJ (p < 0.001) (Fig. 5H). While TRAV41 was most enriched in KIR3DL3+ αβ T cells, the effect was still significant with this gene removed (Fig. 5H, p = 0.02). Agonist-selected CD8αα+ tissue-resident T cells fail to downregulate *TRGC,* express Hobit, a transcription factor that coordinates tissue-residency and innate T cell phenotypes, and Helios, which can be induced upon strong TCR stimulation ^63,67^. We observed upregulation of *TRGC2, ZNF683* (Hobit), and *IKZF2* (Helios) in KIR3DL3+ αβ T cells (Fig. 5I). Additionally, KIR3DL3+CD8+ T cells of peripheral blood and jejunum exhibited a reduced ratio of CDβ:CD8α, which has been described as an indicator of CD8αα expression in humans ^68^ (Fig. 5J). Together, these results suggest that KIR3DL3+ CD8+ T cells may represent a population of agonist-selected T cells.

### KIR3DL3 polymorphism having unique population genetic patterns does not impact ligand binding

KIR polymorphism can drastically alter receptor expression, ligand recognition, and signaling function ^40,48,69–74^. As the KIR3DL3-HHLA2 interaction has potential as an immunotherapeutic target ^23^, the identification of functional KIR3DL3 polymorphism could inform patient selection and expected responses to treatment. We therefore aimed to test a panel of naturally occurring KIR3DL3 variants for their ability to alter KIR3DL3 function or expression.

To confirm that KIR3DL3 is an inhibitory receptor, we transduced the NKL cell line with KIR3DL3*003 containing a C-terminal FLAG tag (NKL-KIR3DL3^FLAG^) and treated cells with the phosphatase inhibitor, pervanadate, to induce tyrosine phosphorylation (Fig. 6A). After pervanadate treatment, we precipitated KIR3DL3^FLAG^ using an anti-FLAG antibody and observed tyrosine phosphorylation and the coimmunoprecipitation of SHP-1 and SHP-2 (Fig. 6B). Recruitment of both SHP-1 and SHP-2 underlie inhibition by KIR3DL1 and other inhibitory KIR ^75–77^, suggesting KIR3DL3 functions with a similar mechanism. Next, we tested whether KIR3DL3 transmits an inhibitory signal using the IIA1.6 mouse B cell line transduced with KIR3DL3*001. We induced B cell activation by crosslinking the BCR with biotinylated Fab anti-IgG and avidin, which induced intracellular Ca^2+^ as evidenced by Indo-1 staining (Fig. 6C). Co-recruitment of KIR3DL3 using biotinylated CH21 significantly reduced Ca^2+^ mobilization (p < 0.05). Therefore, KIR3DL3 can transmit an inhibitory signal, likely through ITIM phosphorylation and SHP recruitment. To confirm that KIR3DL3 can directly recognize the HHLA2 ligand, we recombinantly expressed both molecules and conducted surface plasmon resonance (SPR) experiments, where the HHLA2 ligand was the analyte and KIR3DL3 coupled to the streptavidin sensor chip. KIR3DL3 bound to HHLA2 with an affinity value (K_D_) of 48.9μM (Fig. 6D), which is an affinity within the range observed for KIR3DL1 interacting with its HLA cIass I ligands ^43^.

**Figure 6:**
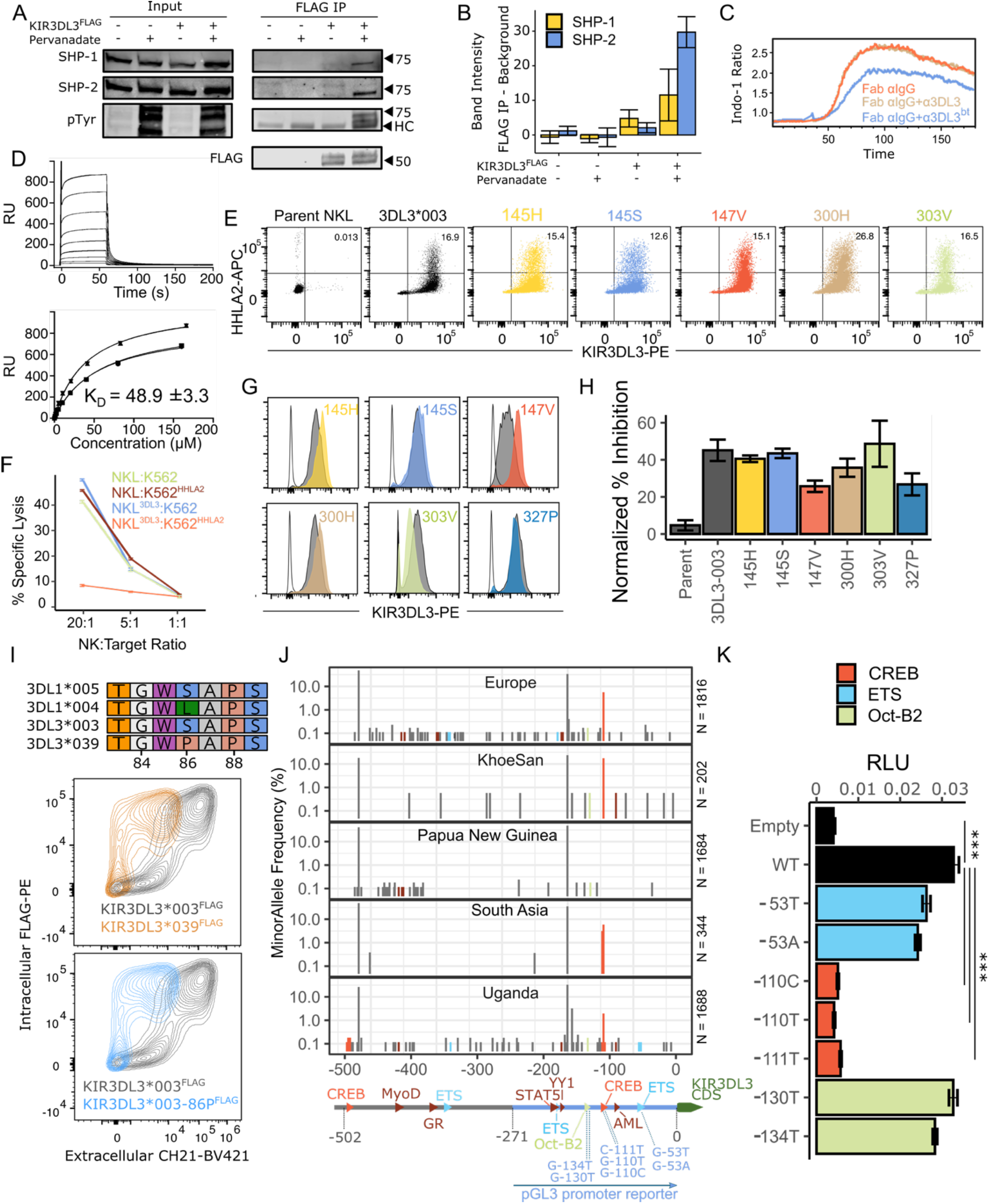
KIR3DL3 is a polymorphic inhibitory receptor with functional promoter polymorphism. (A) Wild-type (WT) NKL or NKL-KIR3DL3*003^FLAG^ were treated with 10 μM pervanadate for 15 minutes, immunoprecipitated with anti-FLAG antibody, separated on a reducing SDS-PAGE gel, and visualized by Western blot through staining with antibodies targeting phosphorylated tyrosines (pTyr), FLAG, SHP-1, SHP-2, or β-tub. HC = heavy chain contamination from IP. (B) SHP-1 and SHP-2 staining was quantified as band intensity (relative to background) following FLAG IP divided by band intensity of input (n = 3). (C) IIA1.6-KIR3DL3 w cells were stained with 2 μM Indo-1 and incubated with Fab anti-IgG alone, with anti-KIR3DL3 (CH21), or with biotinylated anti-KIR3DL3. BCR activation was induced by avidin crosslinking and the 405 nm/485 nm ratio was measured as an indicator of intracellular Ca^2+^. (D) Interactions between HHLA2 and KIR3DL3 were determined using surface plasmon resonance. The equilibrium dissociation constant (K_D_) was determined by flowing serial dilutions of HHLA2 over immobilized KIR3DL3. Top: Sensorgrams for serial dilutions of HHLA2 interacting with KIR3DL3. Bottom: K_D_ was determined by fitting a single site binding model to data from two independent experiments with two replicates each (RU=response units). (E,G,H) We considered the KIR3LD3*003 allotype alongside 6 other variants that occur at naturally selected residues (145H, 145S, 147V, 300H, (extracellular domain) 303V (transmembrane domain), and 327P (cytoplasmic tail)) ^21^. (E) Parental NKLs and the 6 KIR3DL3 variants were co-incubated with 100 μg/mL HHLA2-Fc. KIR3DL3 and HHLA2 were stained with PE and APC conjugated antibodies, respectively, and analyzed by flow cytometry. (F) The indicated NKL and target cell lines were incubated at varying E:T ratios for 5 hours and lysis of K562 analyzed by flow cytometry. (G) Flow cytometry histograms show KIR3DL3-PE staining of parental NKL (empty), control NKL-KIR3DL3*003 line (grey), and the NKL-KIR3DL3 line with the indicated single residue change from KIR3DL3*003. (H) The seven KIR3DL3+ lines were incubated with 221 or 221-HHLA2 cells at a 20:1 ratio for 5 hours. Plotted is the percent inhibition by HHLA2 on NKL cytotoxicity (n=3). (I) Polymorphism at residue 86 defines KIR3DL3*039 and the poorly expressed KIR3DL1*004 allele. 293T cells were transfected with either KIR3DL3*003^FLAG^, KIR3DL3*039^FLAG^, or KIR3DL3*003^FLAG^ with a proline at position 86. Extracellular and intracellular KIR3DL3 were determined by fluorescently labeled CH21 and anti-FLAG, respectively. (J) KIR region DNA was sequenced from individuals from 5 human populations and KIR3DL3 promoter variants identified. Plotted are the minor allele frequency of SNPs within each population, for the 500 bp upstream of the first codon (0). Grey=outside of known TF binding sites; Orange=CREB site; Blue=ETS site; Green=Oct-B2 site; Red=Other untested TF sites. Exact positions mutated within the pGL3 promoter reporter are indicated. (K) Empty pGL3 or pGL3 driven by 8 proximal promoter variants of KIR3DL3, alongside renilla-expressing pRL-CMV, were transfected into 293T cells. Plotted is the ratio of firefly to renilla luciferase as relative light units (RLU). Significance assessed by linear mixed models. *p<0.05, **p<0.01, ***p<0.001

Because functional variant sites are expected to have been the target of natural selection ^78,79^, we focused on polymorphism previously identified at residues with signatures of diversifying natural selection ^21^. We created NKL lines expressing KIR3DL3 with the single residue changes in the D1 domain (R145H, R145S, I147V), transmembrane domain (Y300H, A303V), and cytoplasmic domain (A327P). Each line expressing a KIR3DL3 variant was retrovirally transduced alongside a control “WT” KIR3DL3*003 allele, which were used in HHLA2-Fc binding assays and cytotoxicity assays. We incubated NKL, NKL-3DL3*003, and each KIR3DL3 variant line (excluding 327P, in the cytoplasmic domain) with HHLA2-Fc and observed KIR3DL3+ cells bound HHLA2-Fc across the variants, with minimal binding to NKL alone (Fig. 6E).

We confirmed previous observations ^23,24^ that HHLA2 expression in target cells specifically inhibits lysis by KIR3DL3+ effector cells (Fig. 6F). We then incubated each transduced NKL line with 221 or 221-HHLA2 target cells, in each case comparing to the matched KIR3DL3*003 control line. The polymorphic residues tested did not significantly alter HHLA2-mediated inhibition (Fig. 6G-H).

### KIR3DL3 polymorphism can alter cell surface expression and proximal promoter activity

In addition to ligand binding, KIR polymorphism can reduce receptor expression at the cell surface. To determine if the KIR3DL3 expression level can be determined by genetic variants, we considered polymorphisms at conserved protein residues and in transcription factor binding sites of the proximal promoter. We noticed that the KIR3DL3*039, *052, *059, and *069 allotypes are defined by a serine to proline change at position 86 (86P). This SNP (rs374399247) is rare in most populations but can be as common as 3.5% in certain African populations ^80^. Position 86 is also variable in KIR3DL1, where a serine to leucine mutation results in reduced cell surface expression of the KIR3DL1*004 allotype ^81^. We tested whether KIR3DL3*039 is poorly expressed as compared to KIR3DL3*003 by transfecting 293T cells with C-terminal (intracellular) FLAG tagged KIR3DL3. While levels of intracellular FLAG were similar between KIR3DL3*003 and KIR3DL3*039, extracellular KIR3DL3 staining with CH21 showed markedly reduced cell surface expression of KIR3DL3*039 (Fig. 6I). Specifically mutating residue 86 of KIR3DL3*003 to a proline reproduced the reduced cell surface expression of KIR3DL3*039 (Fig. 6I), suggesting polymorphisms at this position have similar effects in KIR3DL3 and KIR3DL1.

The intermediate and proximal promoter regions have been characterized for KIR3DL3 ^82,83^. Promoter polymorphism in other KIR genes can determine protein expression levels ^84–86^. Therefore, we next considered polymorphic nucleotides in transcription factor bindings sites (TFbs) of the KIR3DL3 promoter ^21,87^. We aligned reads from targeted DNA sequencing of the KIR region to the hg38 genome and identified single nucleotide polymorphisms using samtools ^88^. We identified an ETS site in Papua New Guineans that is triallelic (G-53A/T) and an Oct-B2 site that is triallelic (G-130T in Papua New Guineans, G-134T in Europeans). However, particularly striking was a CREB site with a polymorphism (G-110C) that is absent from some populations but has risen to moderate frequency in others (e.g. 16.5% in KhoeSan) (Fig. 6J). This site is quadrallelic world-wide, although A and T are rare (<1%). Additionally, the nucleotide directly upstream is polymorphic and has the second greatest frequency (3.2% in South Asia) of the TFbs positions (C-111T). These variants are completely unlinked and therefore segregating independently in the human population.

To test whether these natural variants have functional impact, we cloned the KIR3DL3 proximal promoter upstream of luciferase and introduced the relevant mutations. We transfected 293T cells and measured luciferase as a proxy for promoter activity. The KIR3DL3 proximal promoter induced expression of luciferase. Mutations in the Oct-B2 site (G-130T, G-134T) did not affect luciferase expression (Fig. 6K). Those (G-53A, G-53T) in the ETS site only modestly reduced expression. Remarkably, those in the CREB site (G-110T, G-110C, C-111T) completely ablated proximal promoter activity (p < 0.001). To rule out off-target mutations from the PCR-based mutagenesis procedure, we independently cloned a proximal promoter from a South Asian individual homozygous for −110C and observed a similar effect (Fig. S6). We also repeated experiments in 221 cells to rule out cell typespecific effects (Fig. S6). We conclude that −110 (C=rs111269305) has unique population genetic patterns and segregates multiple mutations with potential to remove proximal promoter activity. These results raise the possibility of differential KIR3DL3 expression across individuals.

## Discussion

The expression profile of KIR3DL3 has remained largely uncharacterized, perhaps in part due to an expectation that it would be primarily expressed in NK cells. Instead, we found KIR3DL3 is expressed by CD8+ T and γδ T cells of PBMC, thymus, lung, and multiple regions of the digestive tract. Intraepithelial lymphocytes are comprised of T cells that express αβ or γδ rearranged TCRs, their functions include maintaining epithelial integrity and responding rapidly to infections ^89,90^. That KIR3DL3 can be expressed in over 50% of γδ IELs suggests an important biological role for this inhibitory receptor in the digestive tract. Indeed, CD28H, an activating receptor that also recognizes the HHLA2 ligand, is expressed on IELs ^91^, and HHLA2 is most highly expressed in the small intestine epithelium ^51^. As such, an interplay with KIR3DL3-HHLA2 likely balances the ever-present CD28H-HHLA2 stimulatory signal. This may raise the activation threshold for IELs, and allow stronger immune responses upon HHLA2 downregulation, akin to missing-self responses mediated by other inhibitory KIR, although further studies are required to confirm this.

We identify several lines of evidence that suggest KIR3DL3 is upregulated alongside or downstream from TCR stimulation. Single cell RNA sequencing indicated a subset of canonical TCR-responsive and NF-κB-regulated genes are upregulated in KIR3DL3+ T cells. KIR3DL3+ T cells upregulate surface expression of CD69, CD27, and CD56, each of which can be upregulated following T cell activation ^92,93^. Surface expression of CD3 is also reduced, which can be observed following TCR stimulation or in agonist-selected CD8αα T cells ^63,94^. Although these observations suggest KIR3DL3+ T cells have been recently stimulated, we note that we did not observe increased KIR3DL3 expression following incubation with anti-CD3/CD28 beads (Fig. 4G). This observation is contrary to a previous report that the frequency of KIR3DL3 expression is increased on T cells following TCR stimulation and long-term culture ^23^. Alongside the exceedingly rare expression of KIR3DL3 in PBMCs across previous datasets spanning diverse disease states (Fig. 1A), these observations suggest that canonical TCR stimulation alone is unlikely sufficient to upregulate KIR3DL3.

KIR3DL3+ T cells in peripheral blood share many attributes with unconventional IELs, which have been described as having a “resting-but-activated” phenotype ^95^. Innate-like CD8αα+ intraepithelial T cells are enriched for auto-reactive T cells, can be hyporesponsive to TCR engagement, are Vδ1-biased, rearrange TCR-α chains early in agonistic thymic selection, and upregulate NK cell inhibitory receptors and TCR-responsive genes ^90,95–97^. The transcriptional profile, cell surface receptor phenotype (CD3^low^CD27^hi^CD69+CD56+), and TCR repertoire of KIR3DL3-expressing T cells reflect these features. KIR3DL3+ cells are hypofunctional, with a compromised metabolic signature, and upregulate genes canonically downstream of TCR stimulation, consistent with a model of anergy induced by chronic TCR stimulation or agonistic selection ^98–100^. Additionally, KIR3DL3 is preferentially upregulated in Vδ1+ T cells or CD8+ TCRαβ T cells with early rearranging TCR-α chains, both of which can have autoreactive properties ^63,101–103^. These patterns are consistent with an unconventional mode of T cell activation associated with KIR3DL3 expression. The similarity between KIR3DL3+ peripheral T cells and IEL populations may suggest a common developmental origin or the eventual recruitment of circulating KIR3DL3+ cells to HHLA2+ tissues, such as the lungs or digestive tract. Alternatively, peripheral KIR3DL3 expression could be upregulated during the early differentiation of “induced” conventional IELs ^90,95^.

There is evolutionary evidence supporting KIR3DL3 upregulation downstream of the TCR. In an amazing example of convergent evolution, separate gene families in humans and mice have radiated and sub-functionalized to perform the same tasks: KIR in humans and Ly49 in mice ^104^. Both KIR and Ly49 have inhibitory and activating members that are highly polymorphic, expressed on NK cells, and mostly recognize MHC class I molecules as ligands ^105^. However, the mouse Ly49E is unique, with striking similarities to the characteristics of human KIR3DL3 that we have described. Both are framework genes encoded on the far edge of their respective gene clusters. Ly49E recognizes the non-MHC urokinase plasminogen activator molecule as a ligand ^106^, is induced downstream of TCR stimulation in peripheral blood ^107^, and is expressed in intestinal intra-epithelial γδ T cells ^107,108^. Ly49E is also found on Vγ3+ thymocytes, Vγ3+ epidermal T cells, and fetal NK cells ^109–111^, and we note the observation of KIR3DL3 expression in multiple skin samples in the SRA (Fig. 1A) and in decidual lymphocytes ^31^. These similarities suggest that convergent evolution of KIR and Ly49 gene families extends beyond missing-self MHC recognition to include a T cell inhibitory receptor with roles in maintaining epithelial homeostasis and possibly in coordinating reproduction.

Guided by patterns of natural selection, we tested whether protein polymorphism in KIR3DL3 could alter HHLA2 recognition or effector responses. For example, the 145RSH and 147IV variants are maintained at equivalent positions in human lineage III KIR and chimpanzee KIR3DL3, respectively ^21^. We did not observe strong differences in HHLA2 binding or effector responses across polymorphic residues. Such strong natural selection may be expected to have more robust consequences, especially given the abundance of functional diversity found in other KIR. Thus, the molecular and biological underpinnings of the population genetic patterns observed for KIR3DL3 remain a mystery. Future work should consider context-dependent functions for KIR3DL3 polymorphism across tissues, during infection, or in determining affinity for undiscovered ligands other than HHLA2.

We identified multiple polymorphic nucleotides in a CREB binding site of the proximal KIR3DL3 promoter that ablate activity of luciferase-based promoter reporters. At face value, this observation could suggest individuals homozygous for (e.g.) −110C would not express KIR3DL3 protein in certain circumstances where the CREB transcription factor is required for expression. In a model where KIR3DL3 is induced as an immune checkpoint alongside CD28H, this could bias CD28H+ lymphocytes toward continued activation. However, in addition to the proximal promoter, KIR3DL3 can be controlled by an intermediate promoter, and the possibility of compensatory mutations upstream of the proximal promoter cannot be presently excluded.

Together, our data suggest a conserved and essential function for KIR3DL3 in activated and innate-like T cells at mucosal sites such as the lung and intestine. A need for its biological function has likely led to the convergent origination of KIR3DL3 and Ly49E, with diversifying natural selection further differentiating the former across human evolution. Future studies on the significance of this receptor in infectious, inflammatory, and malignant disease may reveal the biology driving natural selection pressures on KIR3DL3 and further inform the utility of KIR3DL3-HHLA2 as an immunotherapeutic target.

## Materials and Methods

### Human Tissues

Healthy donor de-identified peripheral blood was obtained by the University of Colorado Clinical and Translation Research Centers (CTRC), part of the Colorado Clinical and Translation Sciences Institute (CCTSI), in sodium heparin tubes and PBMC isolated by Ficoll gradient (Cytiva). Additional healthy control samples, plateletpheresis leukoreduction filter (LRS chambers) were purchased from Vitalant Blood Center (Denver, CO, USA). Thymus tissues were obtained and processed within 1 hour after extraction from infants undergoing corrective surgeries for congenital heart disease. De-identified human lungs were procured from deceased organ donors by either Donor Alliance (Denver, CO, USA) or the International Institute for the Advancement of Medicine (Edison, NJ, USA). Donors were nonsmokers who had no history of lung disease, including no emphysema or other smoking-related lung disease, and who died of non-pulmonary causes. All donors were ventilated endotracheally for three days or less prior to organ donation. Lung tissues were processed immediately upon arrival and always within 24 hours of death. The main pulmonary artery was perfused with phosphate buffered saline (PBS) until the venous output was clear. Subjects had no history of lung disease, including no emphysema or other smoking-related lung disease. Healthy sections of jejunum, duodenum, and colon tissues were obtained from organ donors and patients undergoing surgical resections and ethical approval was granted by the Colorado Multiple Institutional Review Board (COMIRB) protocol #21-4748. The donor metadata are summarized in Table S3.

For thymus, tissue was placed in complete RPMI 1640 media (Gibco, #22400-071) (10% heat-inactivated fetal bovine serum (FBS, Sigma-Aldrich), 1% non-essential amino acids (Sigma-Aldrich), 1% Sodium Pyruvate (Sigma-Aldrich), 1X GlutaMAX (Gibco), 1% Penicillin/Streptomycin (Gibco), and 1X 2-mercaptoethanol (BME, Sigma-Aldrich)), cut into small pieces, and gently pressed with the back of a 10 ml syringe to release thymocytes. Thymocytes were then isolated by Ficoll gradient. Lung tissues (10-25 grams/donor) were minced to a liquid consistency, prior to dilution with digestion mix (4.0 mg/mL Collagenase D (Sigma-Aldrich) and 0.6 mg/mL DNAse I (Sigma-Aldrich), diluted in RPMI), then transferred to gentleMACS C tubes. Lymphocytes were dissociated using gentleMACS program m_lung_01_02 five times followed by m_liver_03_01 once, incubated while shaking at 37°C for 1 hour, then one round of the m_liver_04_01 program. Tissue debris was filtered using a metal cell strainer and red blood cells were lysed with ACK lysis buffer. Jejunum, duodenum, and colon samples were minced into IEL dissociation buffer (PBS, 10 mM DTT, 5 mM EDTA, 10 mM HEPES, 5% FBS) and incubated at 37°C while shaking for 30 min. IEL dissociation buffer was replaced once and this process repeated. Debris was filtered and red blood cells lysed with ACK lysis buffer. Lymphocytes from all sources were washed twice in complete media before cell surface staining or cryopreservation in FBS and 10% DMSO.

### Identification of KIR3DL3 reads in Short Read Archive samples

To identify RNA sequencing datasets containing KIR3DL3 transcripts, the first 1,000,000 fastq reads were downloaded from each of the human transcriptomic studies with PolyA selection that were publicly available in the Short Read Archive (SRA). For each downloaded sample (N=360,787), the raw sequencing text data was searched for CTTGCAGGGACCTACAGATGCTTTGGTTCTGTCACTCACTTACCCTATGAGTTGTC, corresponding to position 565-620 in the KIR3DL3 CDS. This sequence is conserved across KIR3DL3 alleles and unique to KIR3DL3 in comparison with other KIR. Traditional mapping methods for the highly polymorphic and homologous KIR gene region would be unable to map reads of KIR3DL3 from RNA transcripts. KIR3DL3+ SRA records were combined according to donor ID, and those that were *ex vivo* or *in vitro* experiments were removed.

### Determination of KIR3DL3 antibody specificity

We tested the binding properties of three anti-KIR3DL3 antibody clones: 1136B (R&D), 26E10 ^24^, and CH21 ^31^. 1136B could appropriately identify cell lines transduced with KIR3DL3, but exclusively recognized pre-apoptotic cells in PBMCs, as evidenced by colocalization of Annexin V and 1136B (Fig. S1A), suggesting this clone is not appropriate for *ex vivo* staining. Additionally, staining of PBMC (Fig. S1B-C) and KIR-transduced cell lines (Fig. S1D-E) with 26E10 showed this antibody reacts with both KIR3DL1 and KIR3DL3, suggesting most 26E10+ cells in peripheral blood are KIR3DL1+KIR3DL3-. Re-analysis of published transcriptome data on sorted 26E10+ PBMCs ^24^ supports this conclusion (Fig. S1F). Analysis of KIR gene expression in scRNA-seq data was assessed using BLAST to assign reads to KIR genes and alleles, keeping only reads that could be uniquely assigned to a KIR locus (Fig. S1G). CH21 staining of transduced cell lines (Fig. S1D-E) and KIR expression in scRNA-seq data support CH21 as a specific monoclonal antibody targeting KIR3DL3 (Fig. S1G).

### Flow cytometry and KIR3DL3 antibody staining and magnetic bead-based enrichment

To enrich for KIR3DL3+ cells, up to 1×10^9^ cells were incubated with CH21 (1:200) in MACS buffer (0.5% BSA, 2mM ETDA, PBS), washed twice, incubated with PE-labelled anti-Ms IgG (1:300), washed twice, then incubated with anti-PE microbeads (1:20) (Miltenyi). Microbead-labelled cells were passed through an LS column (Miltenyi) and the enriched fraction stained with the full panel of antibodies alongside the pre-enriched cell population. See Table S4 for all antibodies used in this study. Intracellular staining for granzymes and perforin was performed with BD Transcription Factor Buffer Set according to the manufacture’s specifications. Unless otherwise indicated, flow cytometry data was collected on a Cytek Aurora flow cytometry system using SpectroFlo software (v3.0). Data were analyzed using CytoExploreR (v1.0.8) and plotted with FlowJo (v10.7.1). All statistics were performed using generalized linear mixed models fit in MCMCglmm ^112^ (v2.33), with statistical code and raw data available on figshare (doi: 10.6084/m9.figshare.20422182).

### Single cell RNA sequencing library preparation

Single cell whole transcriptomes and TCR sequencing libraries were prepared using the BD Rhapsody Single-Cell Analysis System (BD Biosciences) according to the manufacturer’s specifications. From 5 PBMC donors, 4 populations were sorted after viability, doublet, B cell (CD19+) and monocyte (CD14+) discrimination: 1. NK cells (CD3-CD56+), 2. CD8+ T cells (CD3+CD8+), 3. γδ T cells (CD3+TCRγδ +), and 4. Enriched KIR3DL3+ (CH21+) cells. Cell subsets for the populations from different donors were labelled simultaneously with the cell surface sorting panel (Table S4) an oligonucleotide-tagged antibody sample tag (BD Biosciences) prior to cell sorting on a FACSAria3 (BD Biosciences) (Fig. S7). Prior to cDNA library preparation for the WTA and VDJ libraries, all cell subsets from the different donors were pooled, up to 12 unique sample tags were combined per library. Libraries were sequenced on an Illumina NovaSeq.

### scRNA sequencing analysis

Quality control, demultiplexing, and read mapping were performed using the BD Rhapsody WTA Analysis Pipeline (v1.0) on the Seven Bridges server. Count and meta data were loaded into Seurat (v4.1.0) for quality filtering of cells based on mitochondrial contamination (<25% of UMI counts) and sequencing breadth (200-4000 expressed genes per cell) ^53^. Normalization, clustering, dimension reduction, and multimodal reference mapping were also performed in Seurat. For the latter, the dataset was mapped to a CITE-seq reference dataset ^53^, transferring over the population labels to the cells. Based on these labels, contaminating CD4+ T and B cells were removed from the dataset. Analyses was focused on NK, CD8+ T, and γδ T cell subsets. Differential gene expression was performed between KIR3DL3+ and KIR3DL3-of the following subsets: naïve CD8+, central memory CD8+, effector memory CD8+, and γδ T cells. The top 10 genes from each of these comparisons are presented in Fig. 4D. VISION (v3.0.0) was used to calculate a Naïve Score and a TCR Activation Score, which is the sum of normalized expression values of a chosen gene set. Genes for the naïve score were those genes that specifically defined (p<0.001) naïve CD8+ T cells in our data. Genes for the TCR activation score were established immediate-early or early TCR-responsive genes with prior evidence of RNA upregulation following TCR stimulation. The fgsea R package (v.1.20.0) was used for gene enrichment analyses ^58^. For VDJ TCR repertoire analyses, frequencies of shared TRAV, TRBV, TRGV, and TRDV genes across the five donors were tested for differences between KIR3DL3+ and KIR3DL3-cells. A mixed effects model in MCMCglmm was used with a fixed effect associated with KIR3DL3 expression and random effects associated with donor, V gene, and the interaction between V genes and KIR3DL3. The latter random effect was used to determine if V gene usage was significantly altered in KIR3DL3+ cells. All data and code required to replicate analyses are available on figshare (doi: 10.6084/m9.figshare.20425128).

### T cell receptor stimulation

One million cells PBMCs were incubated in 24 well plates with 30 IU/mL IL-2 (NCI) and with or without 25 μL of Human T-Activator CD3/CD28 dynabeads (ThermoFisher). Following 48 hours, beads were removed by magnetic separation. At 7 days cells were stained with viability dye-efluor506, CH21, anti-Ms IgG PE, and fluorescent-conjugated antibodies, anti-CD3, CD56, CD14, CD19, TCRγδ, Vδ1, and Vδ2 (Table S4). Samples were acquired on the Cytek Aurora to determine the proportion of KIR3DL3+ cells and total T cells.

### Cell culture

NKL, K562, and 721.221 cells were cultured in RPMI 1640 media (Gibco, #22400-071) and the Phoenix Ampho packaging line in DMEM (Gibco, #11995-065), at 37°C with 5% CO_2_. All cell culture media was supplemented with 10% FBS (Sigma), 1X GlutaMAX (Gibco), and 1% Penicillin/Streptomycin. NKLs were additionally supplemented with 200 IU/mL of IL-2 (NCI).

### Cloning

The cDNAs for C-terminally FLAG-tagged *KIR3DL3*003, KIR3DL3*039, KIR3DL3*003-86P, KIR3DL2*001, KIR3DL1*015* and *HHLA2* were synthesized by Integrated DNA Technologies. KIR cDNAs included 5’ BamHI and 3’ NotI sites and were cloned into the pMXs-IRES-puro retroviral expression vector (CellBio Labs). HHLA2 cDNA included 5’ EcoRI and 3’ NotI sites and was cloned into the pBMN-IRES-GFP vector (Addgene #1736). PCR (Q5 enzyme; New England Biosciences) with overlapping primers (Table S5) was used to introduce mutations into pMXs-KIR3DL3*003^FLAG^-IRES-puro. PCR products were DpnI-treated and transformed into OneShot Top10 competent cells (ThermoFisher).

The KIR3DL3 proximal promoter (271 bp upstream of the first codon) was amplified from a single South Asian individual ^79^, using primers (Table S5) that introduced a 5’ KpnI site and a 3’ XhoI site and cloned into the pGL3 firefly luciferase vector (Promega). Site directed mutagenesis was performed as described above, with overlapping primers introducing each mutation of interest (Table S5).

### Creation of KIR+ NKL and HHLA2+ K562/721.221 cell lines

Phoenix Ampho cells (0.5 x 10^6^) were plated in a 60mm dish. At 70-80% confluency, Ampho cells were washed with OptiMEM and transfected with 8 ug of retroviral vector using Lipofectamine 3000 (16 μL) and PLUS reagent (40 μL) following manufacturer’s instructions (Invitrogen) ^113^, incubated for 3 hours in OptiMEM, then overnight in DMEM media. At 24 hours following transfection, complete media was replaced with OptiMEM. PLUS reagent and Lipofectamine were frozen overnight before use ^114^. Virus-containing OptiMEM media was harvested at 24 hour intervals and replaced with fresh media for another 3 days. At 48 hours post-transfection, virus supernatant was filtered through a 0.44 um PVDF membrane (Millipore), incubated with Lipofectamine 3000 and PLUS reagent as above, and used to infect 1×10^6^ NKL, K562, or 721.221 cells. Retroviral transduction was performed in 6 well plates. Cells and retrovirus were spun at 700 x g for 30 minutes, incubated at 37°C for 3 hours, followed by centrifugation at 700 x g for 30 minutes. 4 mL of RPMI media/10% FBS + 1000 IU/mL IL-2 was added to the cells in supernatant and incubated overnight at 37°C. Media was replaced with 4 mL RPMI + 4 mL of virus supernatant for the following two days (72 and 96 hours following transfection). Upon confluency, K562 and 721.221 cell lines were cell sorted based on HHLA2 expression. NKL transduction efficiency remained low and were treated for one week with 2 μg/mL puromycin to enrich for KIR-transduced cells, prior to sorting on a BioRad S3e sorter (BioRad).

#### Protein production

KIR3DL3 and HHLA2 were expressed in Expi293™ cells (Gibco) and Hi-5 cells cells (Invitrogen), respectively. The extracellular domain of KIR3DL3*001 (residue 22-320) was cloned into pHLsec ^115^ encoding a C-terminal BirA tag and hexa-histidine tag. Following transfection of Expi293 cells with polyethyleneimine (PEI) ^116^, KIR3DL3 was expressed for 7 days at 37 °C, 120 rpm, 8% CO_2_. Culture supernatant containing KIR3DL3 was dialysed against TBS buffer (10 mM Tris pH 8.0, 300 mM NaCl) and purified using Ni-NTA immobilised metal affinity chromatography (IMAC) followed by size exclusion chromatography (S200 16/60 column; Cytiva). For SPR measurements, purified KIR3DL3 was biotinylated using BirA enzyme and excess biotin removed by size exclusion chromatography (S200 10/30 column; Cytiva). HHLA2 (residue 30-344) was cloned into a modified baculoviral pFastBac-expression vector (Invitrogen) containing a GP64 leader sequence and a C-terminal hexa-histidine tag. HHLA2 was expressed using the Bac-to-Bac™ baculovirus expression system (Invitrogen) according to manufacturer’s instructions. Following infection of Hi-5 cells, HHLA2-producing cells were cultured for 2 days at 21 °C, 120 rpm. Culture supernatant was buffer exchanged into TBS using a tangential flow filtration system (Millipore) and purified via IMAC followed by S200 size exclusion chromatography in TBS. Proteins were concentrated using Amicon centrifugal concentrators (Millipore) and purity was assessed by SDS-PAGE (typically > 90%).

#### Surface plasmon resonance measurements

SPR measurements were performed on a Biacore T200 instrument (Cytiva) using a streptavidin (SA) chip. All experiments were performed in HBS buffer (10mM HEPES pH7.5, 300 mM NaCl, 0.005% P20) at a flow rate of 5 μl/min. Biotinylated KIR3DL3 was immobilized to a surface loading of ~2400 response units (RU) and residual biotin binding sites were saturated by injection of 1 mg/ml biotin. Decreasing concentrations of HHLA2 were passed over the chip and recorded response signal was corrected using an empty flow cell as background control. Equilibrium dissociation constants were determined by fitting a single-site binding model to the data. Data analysis was performed using GraphPad Prism version 7.0 (GraphPad).

### Immunoprecipitation and Western Blotting

Either 2×10^7^ or 5×10^7^ (for co-immunoprecipitation) cells were lysed on ice in 1 mL of lysis buffer (30 mM HEPES, 150 mM NaCl, 2 mM Mg(OAc), and 1% NP-40 supplemented with complete protease inhibitor (Roche), 1 mM Na_3_VO_4_, 1 mM PMSF, and 1 mM NaF). Lysate was passed through a 25G needle 20 times, centrifuged at 16000 x g for 30 minutes, incubated overnight with 3 μg of mouse anti-FLAG antibody (clone M2; Sigma), and then for 10 minutes with 2 μg of rat anti-Ms IgG (clone P3.6.2.B.1; eBioscience) to aid immunocomplex formation. Lysate was then incubated for 4 hours with 25 μL of protein G Sepharose 4B beads (ThermoFisher) per 2×10^7^ input cells. Beads and immunocomplexes were spun at 1000 x g for 5 minutes, washed three times in unsupplemented lysis buffer, resuspended in Laemmli buffer, and boiled for 10 minutes prior to separation with SDS-PAGE.

Proteins were separated on polyacrylamide gels in denaturing conditions then transferred to nitrocellulose membranes. Membranes were blocked in 5% BSA in TBST and incubated overnight with primary antibodies targeting the following proteins: FLAG (M2; Sigma; 1:1000), phosphorylated tyrosine (P-Y-1000 MultiMab; Cell Signaling Technology; 1:2000), β-actin (AC-15; Sigma; 1:1000), SHP-1 (C14H6; Cell Signaling Technology; 1:2000), or SHP-2 (B-1; Santa Cruz Biotechnology; 1:1000). Membranes were washed five times in TBST, blocked in TBST + 5% milk, incubated with IRDYE Goat anti-Rb 680 (926-68071; 1:10000) or anti-Ms 800 (926-32210; 1:10000) secondary antibodies, and imaged with a LICOR imaging system.

Band intensity and background was quantified with ImageJ. Background intensity was subtracted from band intensity and used as input for a linear mixed model using MCMCglmm (v2.32) ^112^, with fixed effects associated with pervanadate treatment, cell line, SHP protein, and their interactions, and a random effect associated with experimental block.

### Calcium mobilization assay

IIa1.6-KIR3DL3 cells were created as described above, except using the Phoenix-Eco packaging line and using pMXs-KIR3DL3*001 ^31^. IIa1.6-KIR3DL3 cells were loaded with 2 μM Indo-1AM (eBioscience) and stained with either 0.5 μg/mL or 0.2 μg/mL biotinylated Fab anti-IgG (Jackson Laboratories) and 20ug/mL of biotinylated anti-KIR3DL3 (clone CH21; gift from Dr Ashley Moffett) and coaggregated with 20ug/mL Avidin (Sigma). Relative intracellular free calcium was measured over time using flow cytometry on a BD LSR Fortessa X-20 instrument and analyzed using FlowJo software (v10.8.1). 20 μg/mL of biotinylated CD22 (clone CY34) was used as a positive control ^117^.

### HHLA2-Fc binding assay

NKL cells (1×10^5^) were washed with PBS, stained with Fixable Viability Dye eFluor 780 (eBioScience; 1:20000), washed with FACS buffer (PBS, 0.5% BSA, 0.1% NaN3), then incubated with an HHLA2-Fc fusion protein (Q9UM44; R&D) at 100 μg/mL in PBS for 30 minutes. Cells were stained with anti-KIR3DL3-PE (clone 1136B; R&D) and anti-HHLA2-APC (clone MA57YW; eBioscience) labelled antibodies and fluorescent intensity measured on a Cytek Northern Lights instrument.

### NKL cytotoxicity assays

NKL effector cells were co-incubated in a U-bottom plate with either 721.221 or K562 target cells at varying effector:target ratios, with a total of 5×10^5^ cells/well resuspended in RPMI media/10% FBS + 1000 IU/mL IL-2 for 5 hours at 37°C. Target cells were labelled with Cell Trace Violet (1:2500; ThermoFisher) prior to co-incubation. After co-incubation, cells were washed with PBS, stained with Fixable Viability Dye eFluor 780, washed with FACS buffer, fixed in 1% PFA, and analyzed by flow cytometry as described above. Specific lysis was calculated as the proportion of lysed cells, corrected for background non-specific death. Specific lysis was analyzed in a linear mixed model using MCMCglmm with fixed effects associated with NKL genotype (i.e. parental, KIR3DL3*003, and variant KIR3DL3), target cell genotype (i.e. WT or HHLA2+), and their interaction, and with random effects associated with each experimental block.

### Luciferase assays

1×10^5^ 721.221 or 293T cells were plated in 24 well plates and transfected with 400 ng of pGL3 and 100 ng of pRL-CMV (Promega). Two days later, cells were lysed, and luciferase activity measured with the Dual Luciferase Kit (Promega). Significance was assessed using a linear model (lm function in R).

### Population genetic analysis of the KIR3DL3 promoter

Sequence reads specific to the KIR region were obtained from multiple studies as described ^8,21,118^. The sequence reads were aligned to GRCh38 using bwa (v0.7.17) ^119^ and promoter variants (863 bp upstream of KIR3DL3 start codon) called using samtools mpileup (v1.7) ^88^.

## Acknowledgements

This work was supported by the following funding sources: National Institutes of Health NIAID 1R56AI155729-01 (to P.J.N). ACS IRG #16-184-56 from the American Cancer Society to the University of Colorado Cancer Center (to L.L). W.H.P is supported by NIH Grant F32 AI161790. J.T. was funded by the European Research Council (ERC) under the European Union’s Horizon 2020 research and innovation programme (grant no. 695551). W.J.J is funded by NIH grants R35HL140039 and R01HL130938. We are grateful to Ashley Moffett for providing CH21 monoclonal antibody, and Xingxing Zang for providing 26E10. We acknowledge the BRB Preclinical Biologics Repository for provision of IL-2, the University of Colorado, Anschutz Medical Campus ImmunoMicro Flow Cytometry Shared Resource, RRID:SCR_021321 and Barbara Davis Center flow cytometry services. We acknowledge the University of Colorado, Anschutz Medical Campus Pathology Shared Resource for coordinating donor tissue samples. JR is supported by an NHMRC Australia Investigator award.

## Supplementary Figures and Tables

**Figure S1:**
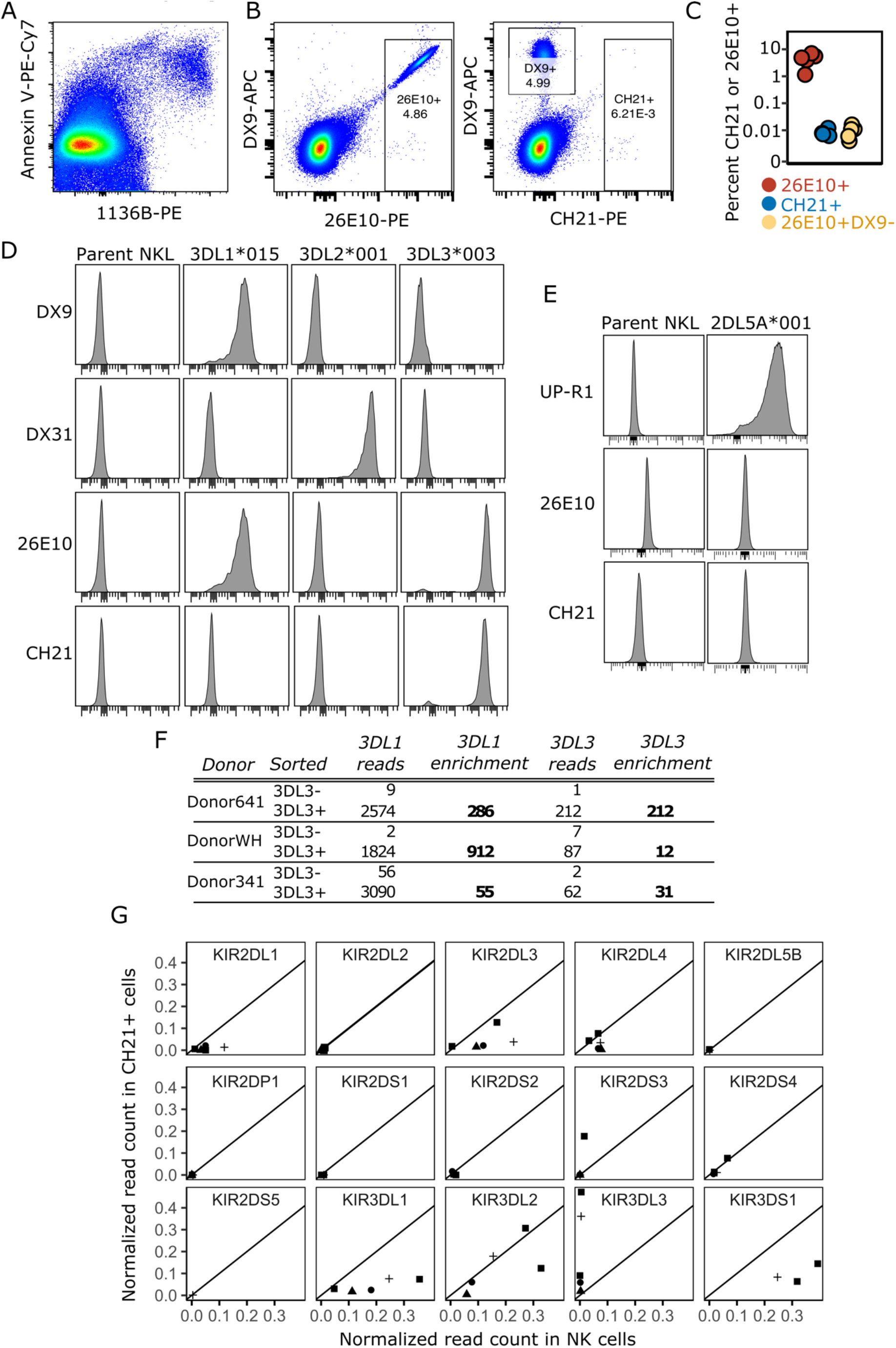
Monoclonal antibody CH21, but not 26E10, is specific for KIR3DL3. (A) Staining of PBMCs with Annexin V and the 1136B antibody, which both detect pre-apoptotic cells. (B) PBMCs from one donor stained for DX9 (anti-KIR3DL1) alongside either 26E10 (left) or CH21 (right). (C) The frequency of 26E10+, CH21+, or 26E10+DX9-subsets were determined within non-B lymphocytes (CD14-CD19-) (n=5 donors). (D) Parental NKLs or NKLs transduced to express cell surface KIR3DL1*015, KIR3DL2*001, or KIR3DL3*003 were stained with DX9 ^120^, DX31 ^121^, 26E10 ^24^, or CH21 ^31^ primary antibodies, followed by PE-labelled anti-Ms IgG secondary antibody. (E) NKL were transduced to express KIR2DL5A*001 and stained with UP-R1 ^122^, 26E10, or CH21. (F) The number of KIR3DL1 and KIR3DL3 reads per sample sequenced in Wei et al, 2021. Enrichment calculated as number of reads in sorted 3DL3+/sorted 3DL3-, as described in Wei et al 2021. (G) KIR specific reads from scRNA sequencing (Fig. 4) were pooled by donor and assigned to KIR genes by performing a BLAST search with each read as a query against the IPD database of KIR alleles, keeping only uniquely assigned reads. This was performed for sorted CH21+ and NK cell (CD3-CD56+) populations, the latter as a measure of background KIR expression. Plotted is the comparison of normalized KIR expression in CH21+ relative to background NK cell expression. KIR enrichment via CH21 evidenced by points above the y=x line. Donors are indicated by shape. Similar results obtained when using sorted T cells as the background population for KIR3DL3.

**Figure S2:**
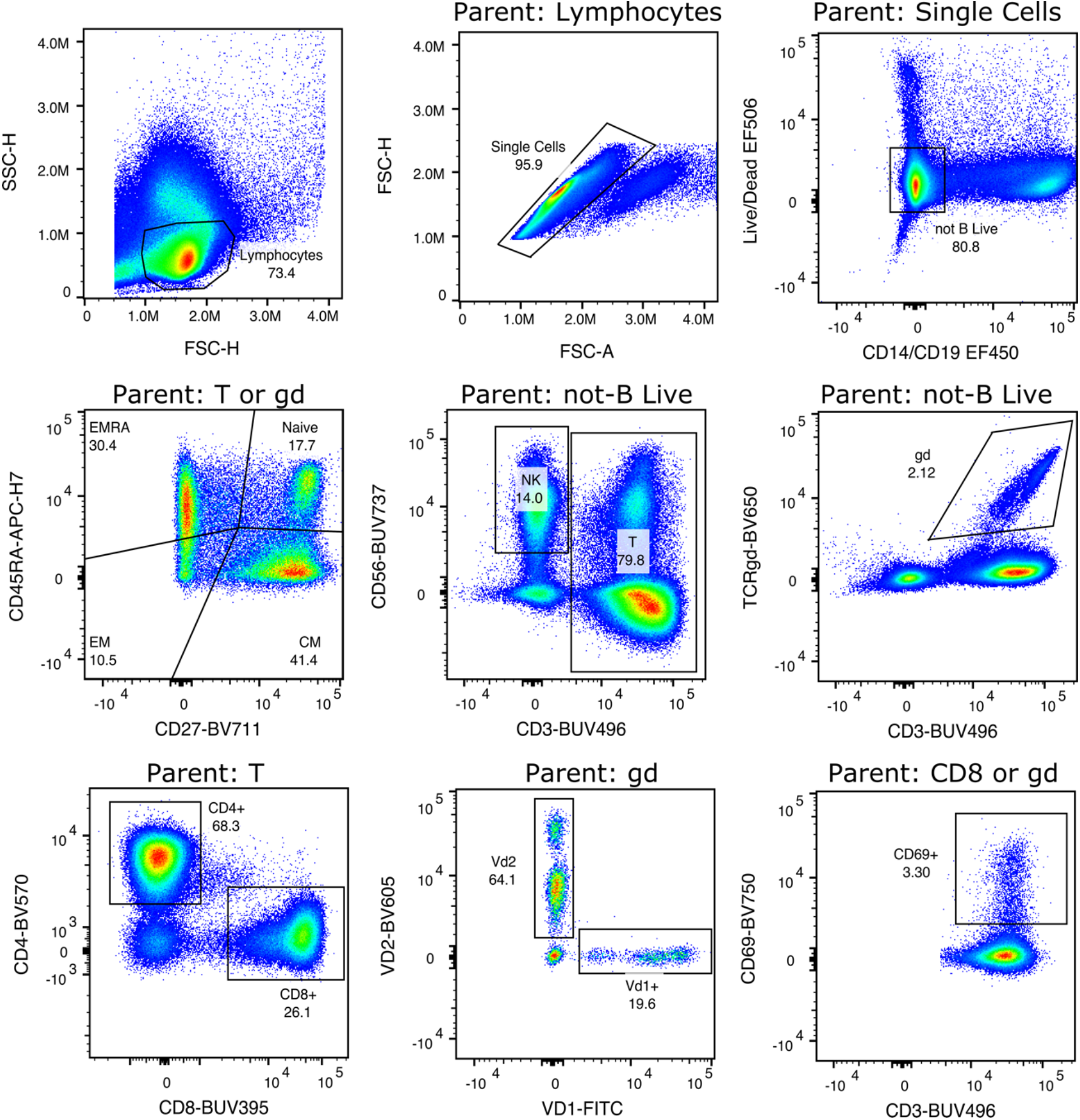
Gating strategy for spectral flow cytometry. Gates are labelled within plots, with parent gates designated above plots.

**Figure S3:**
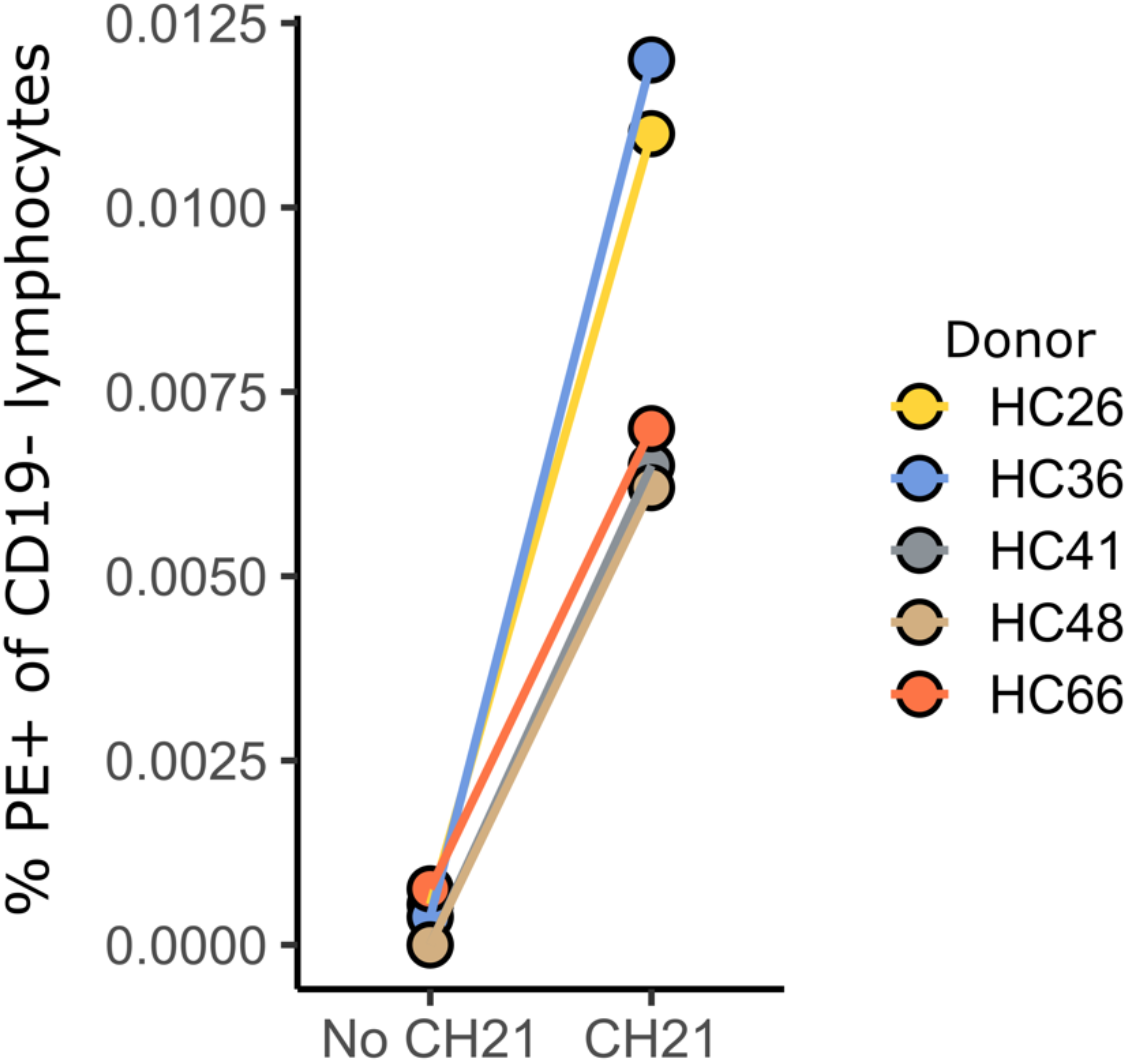
CH21 identifies a distinct population of cells in peripheral blood. PBMCs from 5 donors were cell surface stained with mouse IgG2a isotype control or CH21 followed by secondary anti-Ms PE.

**Figure S4:**
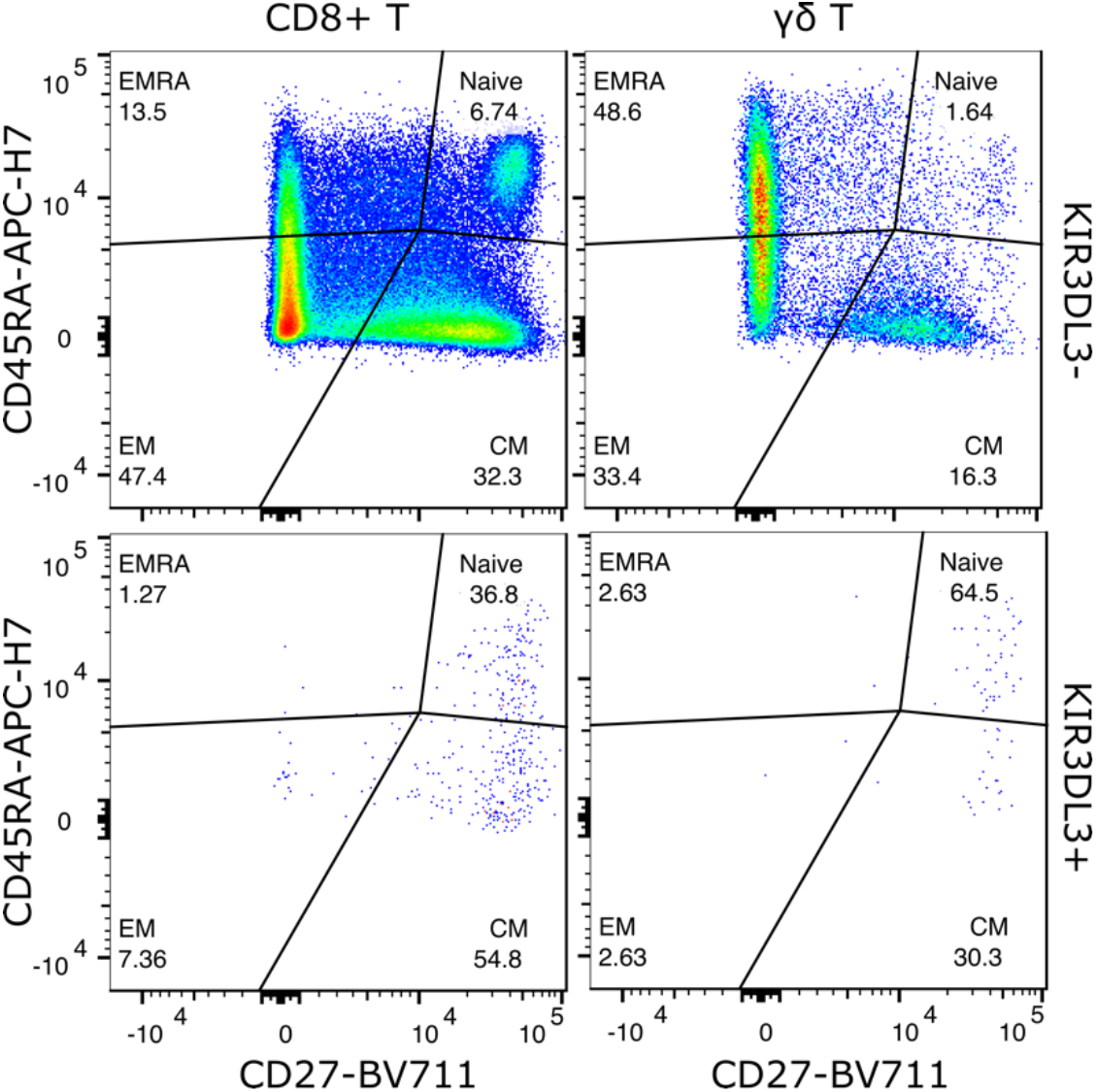
Example of CD45RA/CD27 staining in a single donor. Naive, EM, CM, and EMRA-like populations were determined on the basis of the CD45RA and CD27 markers. Distribution of these subsets are shown for CD8+ T and γδ T cells that are either KIR3DL3- or KIR3DL3+.

**Figure S5:**
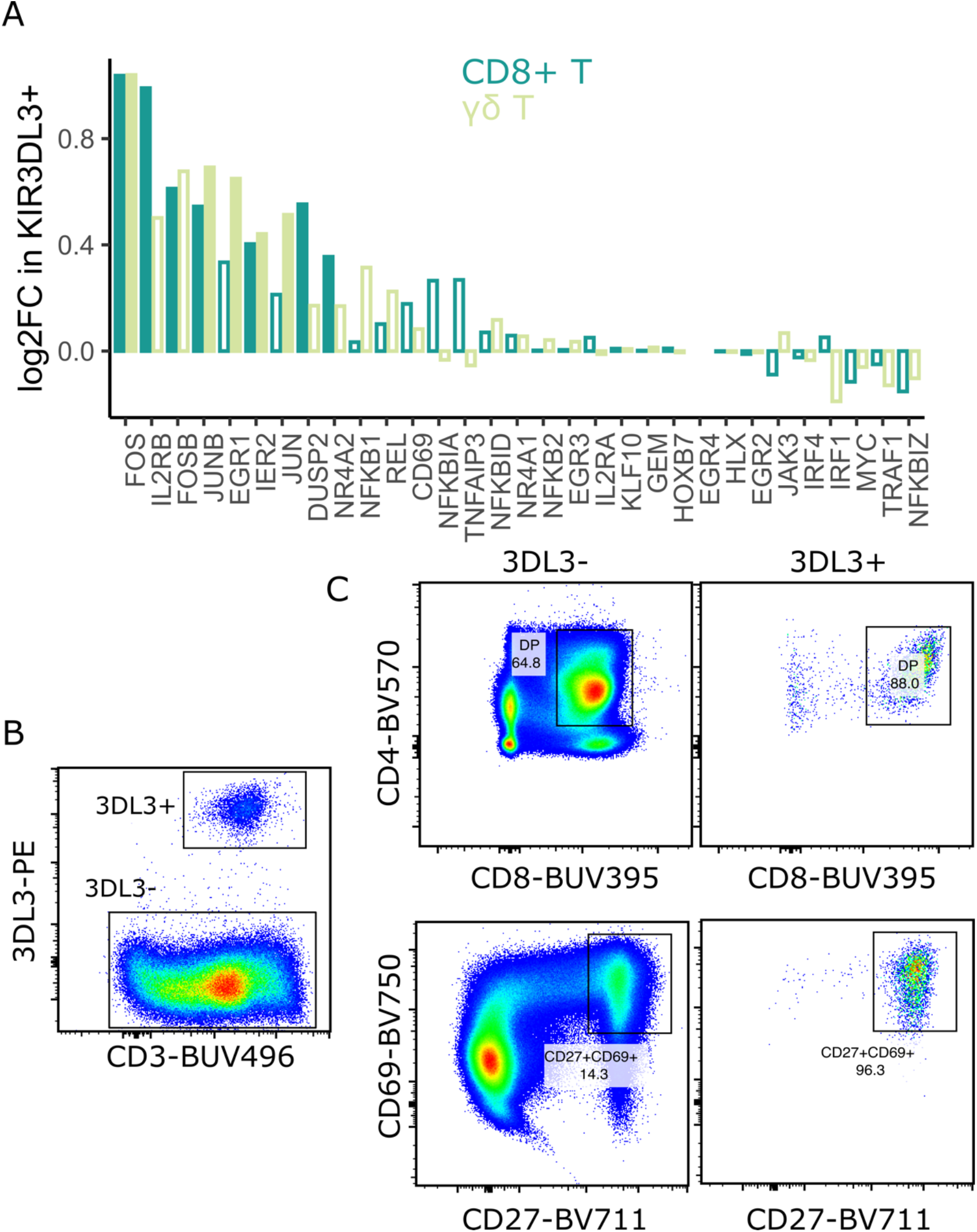
KIR3DL3+ cells co-express TCR-responsive genes. (A) Log2 fold changes of expression in KIR3DL3+ cells are plotted for TCR-responsive genes that were used to calculate the TCR Activation Score with VISION. Closed bars indicate statistical significance at p<0.05 after multiple testing. (B) Thymocytes were enriched for KIR3DL3 by cell surface staining with CH21 followed by secondary anti-Ms PE and anti-PE magnetic microbeads and flow cytometry. Shown gated are either KIR3DL3+ or KIR3DL3-cells of viable CD14-CD19-cells. (C) Cell surface expression of thymocytes stained for CD8, CD4, CD69, and CD27. Gated frequencies are shown as a proportion of either all CD14-CD19-live cells or KIR3DL3+CD14-CD19-live cells.

**Figure S6:**
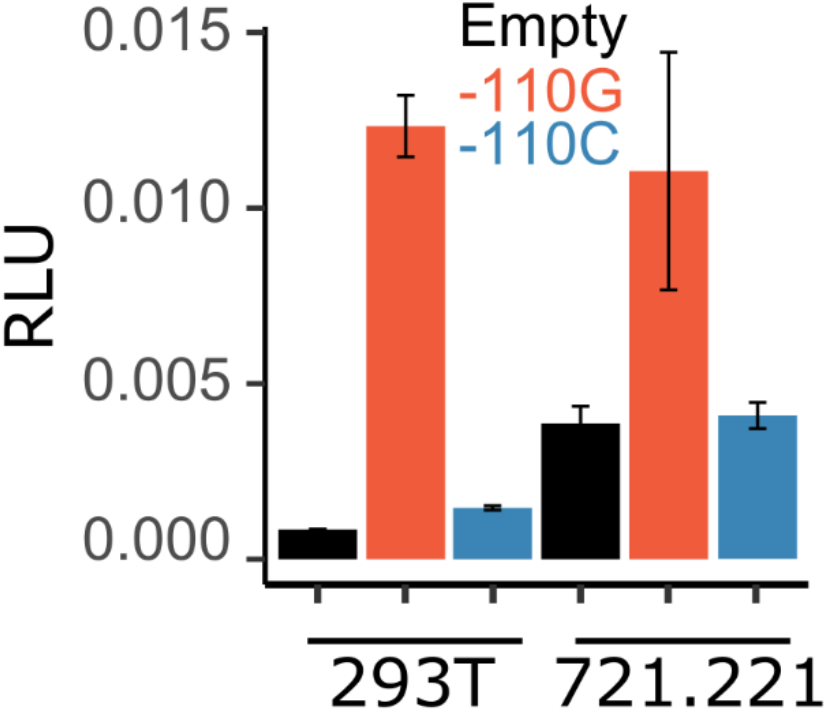
Polymorphism variant −110C reduces KIR3DL3 promoter activity. The KIR3DL3 promoter was cloned into the pGL3 vector from two homozygous individuals having promoters, distinguished only by the −110G/C dimorphism. 293T or 721.221 cells were transfected with pGL3 reporters alongside control Renilla-expressing pRL-CMV. Plotted is the ratio of Firefly to Renilla luciferase as relative light units (RLU). N = 3 per condition.

**Figure S7:**
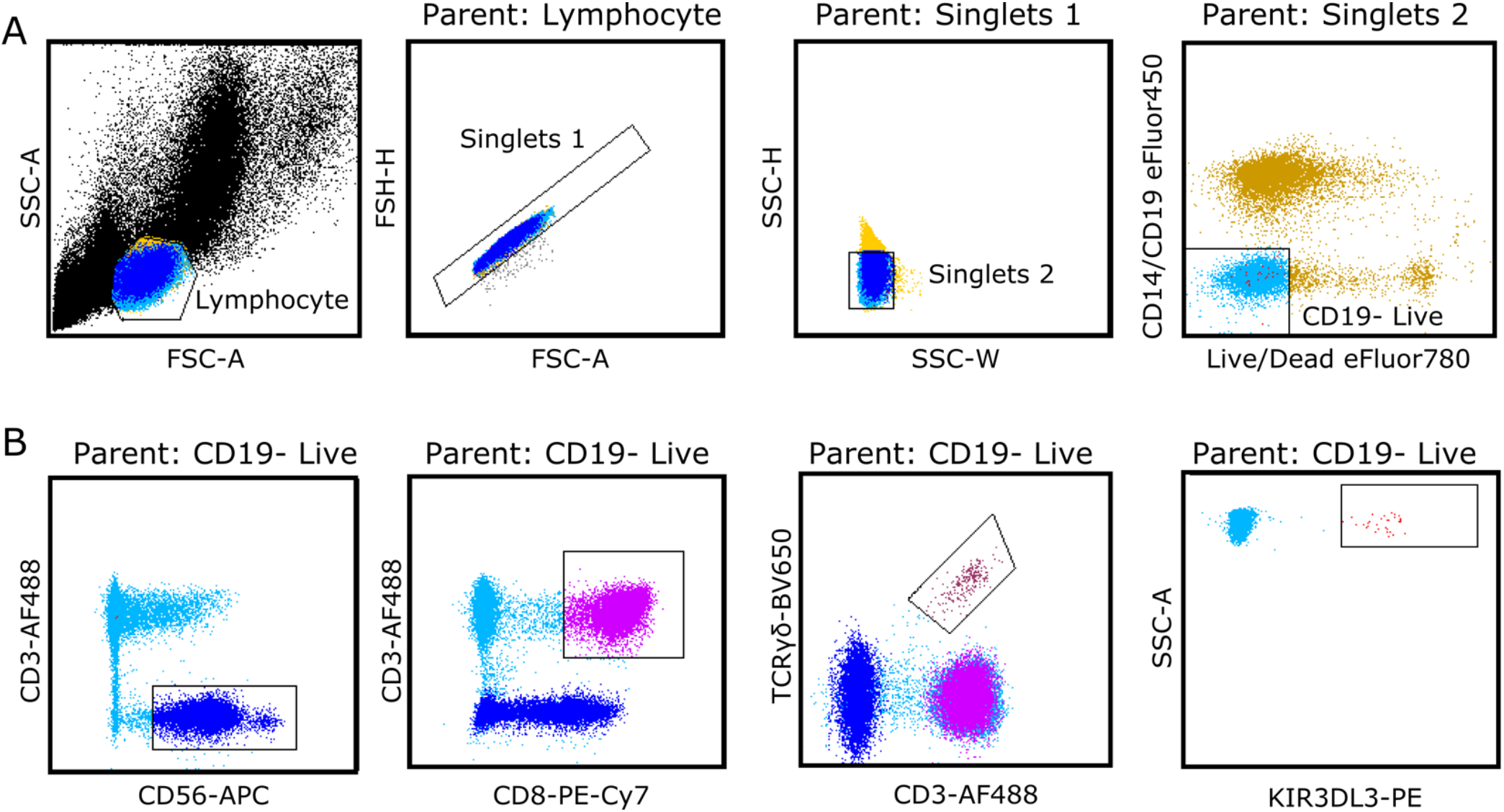
Sorting gating strategy for single cell RNA sequencing. (A) Shows gates set to identify live, CD19-singlet lymphocytes, from which all sequenced populations were sorted. (B) Shows gates used to sort NK cells, CD8+ T cells, γδ T cells, and KIR3DL3+ cells.

Table S1: Meta-data associated with KIR3DL3+ SRA samples.

Table S2: Differentially expressed genes in KIR3DL3+ cells.

Table S3: Blood and tissue donor demographics.

Table S4: Antibodies used for flow cytometry.

Table S5: Primers used for site-directed mutagenesis of KIR3DL3 CDS and promoter constructs.

## Notes

### Competing Interest Statement

The authors have declared no competing interest.

